# Toward navigating emotional states using real-time representational similarity analysis fMRI neurofeedback - a feasibility study

**DOI:** 10.1101/2025.09.25.678461

**Authors:** Xuelei Wang, Assunta Ciarlo, Michael Lührs, Alexander Atanasyan, David Böken, Jürgen Roßmann, Michael Schluse, Maren Jäger, Marisa Nordt, Fengyu Cong, Klaus Mathiak, David E.J. Linden, Rainer Goebel, David M. A. Mehler, Jana Zweerings

## Abstract

Real-time functional magnetic resonance imaging neurofeedback (rt-fMRI-NF) is a promising non-invasive brain-computer-interface (BCI) technique for enhancing self-regulation of affective states in the brain. However, conventional univariate rt-fMRI-NF approaches are limited in their ability to distinguish neural patterns of distinct emotions that involve overlapping brain regions. In this study, we applied an rt-fMRI semantic neurofeedback (rt-fMRI-sNF) paradigm, incorporating real-time representational similarity analysis (rt-RSA) to enable navigation between emotional states. Four emotional patterns were first derived from functional localizer runs, each designed to evoke a specific emotion, and then applied as target patterns during neurofeedback. Using an RSA-informed circular semantic map (CSM), participants received real-time visual feedback indicating both the similarity and intensity of their current brain activity relative to target emotional patterns. Participants were instructed to use mental imagery to shift their brain activity toward the specific target pattern and increase its intensity. Twenty-four healthy participants completed the localizer runs, and two consecutive neurofeedback runs in the same session. Ten participants successfully engaged with both the similarity and intensity components of the CSM, showing effective modulations of their mental states. Analyses of the localizer runs revealed overlapping regional activations across emotions and demonstrated that RSA outperformed univariate analysis in distinguishing between them. For the neurofeedback runs, linear mixed-effects model (LMM) analyses across multiple performance metrics indicated consistent within-run improvements and higher initial performance in the second run, while significant between-run learning effects emerged only in exploratory models with quadratic time terms. A block-wise comparison also showed significantly higher performance at the end of each run compared to the beginning based on the intensity metric. These findings support the usability of RSA in differentiating multiple emotional states and demonstrate the feasibility of the rt-fMRI-sNF paradigm for emotion regulation.

## 1. INTRODUCTION

Real-time functional magnetic resonance imaging neurofeedback (rt-fMRI-NF) has emerged as a promising technique for enabling individuals to learn self-regulating their own brain activity. Among its various applications, emotion regulation is one of the most extensively studied domains (Linhartová et al., 2019). A growing body of research supports the feasibility of rt-fMRI-NF in modulating affective states (Lorenzetti et al., 2018; Weiss et al., 2022; Zhu et al., 2019) and demonstrates its clinical promise in modulating brain activity (Tschentscher et al., 2024) and symptoms in psychiatric conditions such as depression (Marquand et al., 2024; Mehler et al., 2018; Young et al., 2017), posttraumatic stress disorder (PTSD) (Voigt et al., 2024; Zweerings et al., 2018, 2020) and attention deficit hyperactivity disorder (ADHD) (Alegria et al., 2017; Lukito et al., 2024). Traditional rt-fMRI-NF paradigms typically employ a univariate analysis approach, where feedback is based on the average activation level within a predefined brain region and often visualized using a thermometer-like display (Cohen Kadosh et al., 2016). However, this approach has limited ability to distinguish between different mental processes that involve the same brain region, which is a major challenge for emotion regulation because previous studies have shown that different emotions often activate overlapping brain regions (Shibata et al., 2016; Todd et al., 2020). In contrast, multivariate approaches use multivoxel pattern analysis to differentiate between distinct neural activation patterns (Liuzzi et al., 2020; Ritchie et al., 2019), enhancing the capacity of rt-fMRI-NF to target and regulate multiple emotional states.

Building on the advantages of multivariate statistics, one promising method that has gained increasing attention in fMRI research is representational similarity analysis (RSA) (Kriegeskorte et al., 2008; Kriegeskorte & Kievit, 2013). Unlike traditional activation-based analyses, RSA quantifies the relationships between mental representations by examining the representational geometry of neural activation patterns, thereby enabling finer differentiation between closely related cognitive or emotional states. Previous applications of RSA to fMRI data investigated a variety of cognitive processes, including action control (Kolasinski et al., 2020), visual object perception (Grill-Spector & Weiner, 2014; Kriegeskorte et al., 2008; Nordt et al., 2019, 2023) and emotion representation across sensory modalities (Kiyokawa & Hayashi, 2024). In the context of rt-fMRI-NF, a promising real-time RSA application to conduct RSA-informed neurofeedback, known as real-time fMRI semantic neurofeedback (rt-fMRI-sNF), has been introduced (Goebel, 2019, 2021) and evaluated (Ciarlo et al., 2022; Russo et al., 2021) using visual mental imagery tasks. These proof-of-concept studies demonstrated that participants were able to use mental imagery strategies to generate and modulate semantic representations of objects corresponding to external objective sensory stimuli. Applying a similar approach to emotion regulation-based neurofeedback would open new avenues for more personalized self-regulation training and potentially clinical applications (Goebel et al., 2024). However, whether it is feasible to effectively adapt rt-fMRI-sNF to mental imagery of affective states remains to be demonstrated.

A key aspect in rt-fMRI-sNF is the format in which visual feedback is presented to neurofeedback users. Traditional thermometer-like displays only have one degree of freedom and only reflect the overall (e.g., mean-field) intensity of activation within a target region. Thereby, they cannot provide additional information about the relationship (e.g., representational distance) between distinct neural/voxel patterns under different experimental conditions. To overcome this limitation, previous rt-fMRI-sNF studies (Ciarlo et al., 2022; Russo et al., 2021) have introduced a two-dimensional semantic map, which is produced by multi-dimensional scaling (MDS) such that the participant’s current mental state is projected as a moving dot. Specifically, the location of the moving dot is determined by the similarity between the participant’s current voxel pattern (e.g., evoked by mental imagery) and a target voxel pattern (e.g., determined by localizer runs). A semantic map hence allows participants to see how closely their brain activity matches a target mental state. To further enhance the visual feedback information, a novel visualization method was proposed to incorporate both pattern similarity and pattern intensity in a so-called circular semantic map (CSM) (Goebel et al., 2024; Lescrauwaet, 2021). The CSM combines multiple thermometers in a circular arrangement—each indicating the activation intensity of a specific neural pattern—with a dynamic arrow showing the direction of change in pattern similarity. This hybrid form of visual feedback captures two critical aspects of neural representation, highlighting its potential usability in rt-fMRI-sNF.

Here, we conducted a feasibility study for emotion regulation based on rt-fMRI-sNF using a CSM for feedback visualization. Participants were instructed to use mental imagery to induce four distinct emotional states. Individual brain patterns were used to calibrate target states for subsequent neurofeedback runs. Real-time feedback was provided through the CSM, and participants were instructed to modulate their mental states accordingly. Our study aimed to investigate (1) the effectiveness of RSA-based multivariate analysis in distinguishing between multiple emotional states evoked through mental imagery obtained during the localizer scans; (2) the feasibility and efficacy of applying the rt-fMRI-sNF paradigm for emotion regulation; and (3) the usability and potential of navigating brain activity via the CSM-based visual feedback.

## 2. METHODS

### 2.1 Participants

A total of 27 healthy individuals (21 females), aged (mean ± standard deviation (SD)) 28.7 ± 8.3 years, with normal or corrected-to-normal vision and without acute severe neurological, psychiatric, or somatic disorders took part in the main study. A separate pilot group of 6 healthy participants was recruited to assess the experimental design and the usability of task instructions. Data from the pilot group was only used for design validation and excluded from all subsequent analyses. Prior to the scanning session, participants provided written informed consent, and the study was approved by the local ethics committee [EK 23-175].

Because this work was designed as a feasibility study, no formal a priori power analysis was conducted; instead, sample size was determined based on practical constraints and the goal of evaluating methodological feasibility.

### 2.2 Experimental paradigm

This feasibility study was preregistered (https://osf.io/5ybuw). Participants were instructed to engage in four emotional states through mental imagery (joyful relaxation, sadness, enthusiasm and anger) during four sequential functional localizer scans, respectively (Figure 1A). The selection of these four emotional states was based on the valence and arousal model (Feldman, 1995) and informed by a previously published investigation (Heffner & FeldmanHall, 2022). Specifically, two positive and two negative emotions with matched arousal levels (high and low) were selected according to the classification probability of a machine learning model on the valence and arousal ratings of different emotions (Heffner & FeldmanHall, 2022). To deliver instruction and feedback information to the participants, a CSM was used (Goebel et al., 2024), which consisted of four thermometers, each corresponding to a specific emotion. Prior to scanning, participants were informed about the experimental procedure and received standardized task instructions. They were then asked to use mental imagery to separately induce and intensify each of the four emotional states based on their own subjective experience, without receiving any specific imagery instructions beforehand. Subsequently, they reported on a scale from 1 to 10 if they managed to induce the emotional state, how familiar they were with each specific emotion, as well as their perceived valence and arousal levels. In addition, participants were asked to indicate where in the body they sense the emotion. Prior to and after the measurement, all participants completed questionnaires assessing their current mood state and their ability to regulate emotions, as measured by the Profile Of Mood States (POMS) (McNair, 1971) and Emotion Regulation Questionnaire (ERQ) (Gross & John, 2003). At the end of each neurofeedback run, participants were asked to report, on a scale from 0 to 10, the extent to which they were able to induce the target emotion, how well they were able to control their brain activity based on the feedback, and what strategies they used. To enhance the documentation of the study design and reporting, we have enclosed the “Consensus on the reporting and experimental design of clinical and cognitive-behavioral neurofeedback studies (CRED-nf checklist)” (Ros et al., 2020) (see Supplementary Materials).

**Figure 1.**
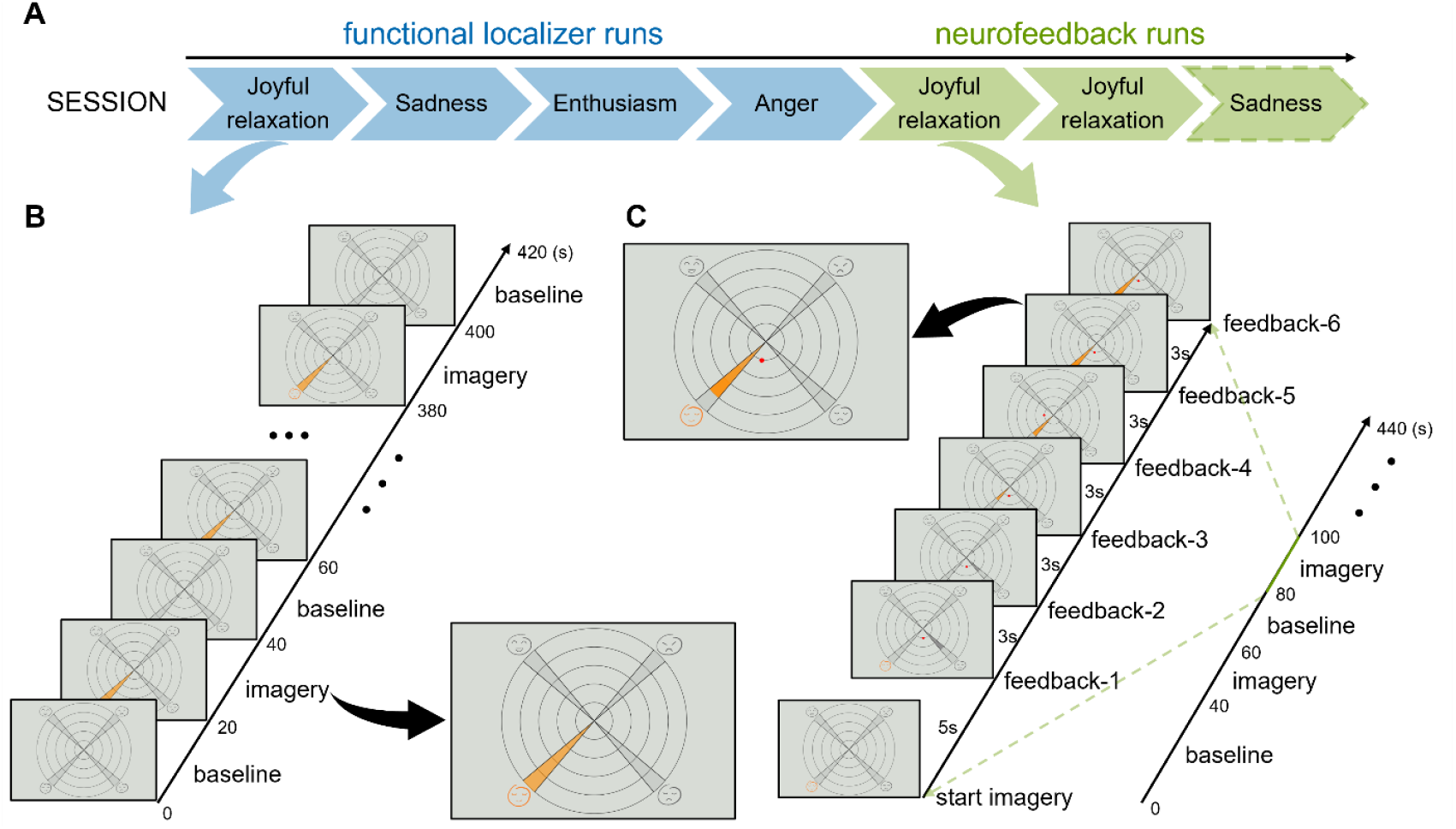
Experimental design. (A) Illustrates the sequence of measurements with four functional localizer runs (blue) and two neurofeedback runs having joyful relaxation as target emotion (green). A third neurofeedback run targeting sadness was optional, and hence, not performed by all participants. (B) Shows the condition sequence within a localizer run for the example of joyful relaxation, with interleaved baseline and imagery blocks. The emotion icon and thermometer (colored in orange during the localizer imagery block) were used as visual cues to prompt participants to perform mental imagery of the corresponding emotion. (C) Shows the condition sequence within a neurofeedback run and one of the mental imagery blocks in detail (exemplified for the emotion of joyful relaxation). During the neurofeedback run, participants were instructed to engage in mental imagery once the emotion icon was highlighted in orange. After 5 seconds, the feedback display was initiated and prompted by a red dot (which indicated the similarity of the current activity pattern with respect to the target pattern) and orange thermometer wedges (which indicated the intensity of the current activity pattern). The three other emotional states that were not targeted during the neurofeedback run were displayed in gray. During each mental imagery block, the feedback display was updated every three seconds, resulting in six trials per imagery block.

#### 2.2.1 Functional localizer runs

The acquisition of an anatomical T1-weighted MRI scan preceded four functional localizer runs (one for each emotion). The functional localizer runs were used to define each participants’ individual target region and to generate the corresponding emotion-specific base patterns. Each run contained 10 emotion imagery blocks interleaved with 11 baseline blocks, both lasting 20 seconds (Figure 1B). Participants were requested to perform the mental imagery task when the icon of a specific emotion and the corresponding thermometer turned orange on the CSM (imagery blocks), and to count backwards from 100 when all emotion icons turned gray (baseline blocks). After completion of the four localizer runs, RSA was used to select voxels with high discernibility between four distinct emotions. The blood-oxygen-level-dependent (BOLD) signal from the selected target voxels served to generate feedback signals throughout the neurofeedback training.

#### 2.2.2 Neurofeedback runs

The experimental design of neurofeedback runs was like that of functional localizer runs, except the initial baseline block was extended to 40 seconds to account for potential preprocessing delays and to stabilize the online statistic computation (Figure 1C). Participants were instructed to imagine one preselected emotion during each of the ten imagery blocks of the neurofeedback runs. The visual feedback provided information to the participants about two key aspects. First, the wedges of the target emotion thermometer were dynamically filled based on the intensity of the current brain activity pattern relative to the corresponding base pattern, extracted during the localizer runs. Second, the current activity pattern was represented as a moving red dot projected onto the CSM. The participants had to adjust their imagery strategy to move the red dot as close as possible to the target emotion, and to increase the target thermometer level.

### 2.3 Data acquisition

MR imaging data was acquired with a 3.0 Tesla Siemens MAGNETOM Prisma whole-body MRI system (Siemens Medical Solutions, Erlangen, Germany), equipped with a 20-channel head coil. A T1-weighted magnetization-prepared rapid gradient-echo (MPRAGE) sequence was acquired for each participant at the beginning of the scanning session and used as anatomical reference (acquisition matrix: 256×256, time of repetition (TR): 2000 ms, echo time (TE): 3.03 ms, 1 mm isotropic voxels, flip-angle: 9°, 176 sagittal slices, duration: ∼4.3 min).

Functional localizer and neurofeedback runs consisted of 420 and 440 volumes, respectively. A multi-band multi-echo echo-planar imaging (EPI) sequence with the following parameters was used for the data acquisition in each functional imaging run: acquisition matrix: 64×64, TR: 1000 ms, TE: [13, 27.8, 42.6, 57.4 ms], flip-angle: 67°, 36 transverse slices per volume that were acquired in interleaved order, inter-slice gap: 0.75 mm, voxel size: 3×3×3 mm, multi-band factor: 3, GRAPPA factor: 2. Partial Fourier acquisition was used with a 7/8 phase-encoding factor to reduce acquisition time.

Data was transferred in real-time from the image reconstruction computer to an analysis workstation for real-time fMRI processing using a direct transmission control protocol/internet protocol (TCP/IP) connection.

### 2.4 Online analysis

Data was processed online using Turbo-BrainVoyager (TBV) version 4.2 (Brain Innovation B.V., Maastricht, The Netherlands) and imported in Python (3.6.12) using the TBV Network Access Plugin and the corresponding interface (https://github.com/expyriment/expyriment-stash/) as part of the open-source Expyriment library 0.10.0 (Krause & Lindemann, 2014). The rt-RSA Python tool, freely available at (https://github.com/assuntaciarlo/rtRSA_GUI), was used for pattern analysis and feedback display.

Based on previous studies (Lindquist et al., 2012; Skottnik & Linden, 2019), we defined a set of emotion-related brain regions and generated a corresponding volume of interest (VOI) in Montreal Neurological Institute (MNI) space. The VOI included cortical regions based on the Schaefer atlas (17-network parcellation, 200 parcels, 1 mm resolution) (Schaefer et al., 2018) and subcortical regions based on the Harvard-Oxford atlas (maximum probability maps, thresholded at 25%, 1 mm resolution) (Desikan et al., 2006). The list of regions in the VOI can be found in Supplementary Table S1.

The online analysis pipeline began by combining multi-echo images using per-volume T2*-weighted sum (Heunis et al., 2021; Mathiak et al., 2002; Weiskopf et al., 2005), followed by real-time data preprocessing and statistical modeling in TBV. The preprocessing steps included motion correction with trilinear interpolation and intra-session alignment (to the first volume of the first functional run) and spatial smoothing with a 6 mm full width at half-maximum (FWHM) Gaussian kernel. The statistical modeling was based on a voxel-wise recursive least squares general linear model (GLM) which was incrementally computed with one predictor for the imagery task, six motion parameters and a linear confound predictor. At the end of each localizer run, TBV performed an offline voxel-wise GLM on the whole time series to extract the stimulus-specific t-map associated with the contrast “task vs rest”. After all the four localizer runs, the rt-RSA tool was used to generate four base patterns by selecting voxels which were related to emotion imagination (univariate selection) and allowed semantic discernibility between emotions (multivariate selection). In the univariate selection, voxels were picked based on their absolute t-values, specifically those whose absolute t-values ranked within the top 60% to 90% of all the voxels within the defined emotion network. The multivariate selection first used a searchlight approach (with a spherical radius of 1 voxel) to generate a four-by-four representational dissimilarity matrix (RDM) for each voxel of the initial mask obtained from the univariate selection. This RDM encoded the dissimilarities between the four emotional patterns obtained from each searchlight, defined as one minus Pearson’s correlation coefficient. Voxels were then selected if their RDMs achieved the top 5% similarity to a predefined model RDM, which treated emotions of opposing valence as maximally dissimilar while disregarding differences in arousal. Specifically, in the model RDM, the dissimilarity is set to 2 between positive and negative emotions (joyful relaxation vs. sadness or anger, and enthusiasm vs. sadness or anger), and 0 between emotions of the same valence (joyful relaxation vs. enthusiasm, and sadness vs. anger). The percentage thresholds and the model RDM were determined based on the initial pilot data (N = 6) and then applied consistently in the main study (N = 27).

During each neurofeedback run, the same preprocessing and statistical modeling steps were performed via TBV. In addition, the current brain activity pattern within the defined target region was correlated with respective base patterns. The correlation results were then used to calculate the location of the red dot and thermometer level (Lescrauwaet, 2021). The red dot was plotted between the thermometer representing the emotion with the highest correlation value and its neighboring thermometer (the one with the next higher correlation). The angular distance ratio of the red dot to the highest correlated emotion thermometer was calculated by the following formula:

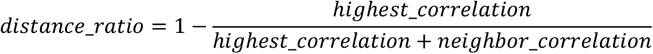

The final location of the red dot was determined by multiplying the distance ratio by the angle between the highest correlated thermometer and its neighboring thermometer. The highest correlated emotion thermometer was filled according to the pattern intensity level and the filling color was orange for the target emotion or gray for non-target emotions. To compute the intensity level, the current activity pattern vector was first projected on the highest correlated base pattern vector. The relative length of the projected vector to the base pattern vector was then referred to as the normalized intensity level. The corresponding thermometer was filled by converting the intensity level into one of five levels, each corresponding to a fixed increment of 0.2.

### 2.5 Offline analysis

The offline MRI data preprocessing was performed using HeuDiConv v1.3.2 (Halchenko et al., 2025) and fMRIPrep v24.1.1 (ignored susceptibility distortion correction and slice timing correction; normalized to MNI152NLin2009cAsym, with 2 mm resolution) (Esteban et al., 2019, 2020). The following analysis, statistical tests and plotting relied on TBV and Python packages, including NumPy 1.19.2 (Harris et al., 2020), Pandas 1.1.5 (McKinney, 2010), SciPy 1.5.2 (Virtanen et al., 2020), Statsmodels 0.12.2 (Seabold & Perktold, 2010), Nilearn 0.9.2 (Abraham et al., 2014), NiBabel 4.0.2 (Brett et al., 2024), Matplotlib 3.3.4 (Hunter, 2007) and Seaborn 0.11.2 (Waskom, 2021). The offline analysis scripts can be found at (https://github.com/xuelei-1021/rsanf).

#### 2.5.1 Neural activity patterns during emotion imagery

We examined brain activity patterns during emotion imagery from both univariate and multivariate perspectives. Univariate analyses assessed traditional task-related BOLD responses using a “task vs. rest” contrast, while multivariate analyses (RSA) characterized distributed voxel patterns and representational dissimilarities between emotions.

##### Univariate activation analyses

The first-level analysis was performed using the FirstLevelModel module in Nilearn (0.9.2), implementing a voxel-wise GLM with high-pass temporal filtering (128s cutoff, cosine basis functions), hemodynamic response function (HRF) modeling from Statistical Parametric Mapping (SPM), time series standardization, spatial smoothing with a 6 mm FWHM Gaussian kernel, and autoregressive (AR(1)) noise modeling. The GLM model was fitted to the time-series data to estimate participant- and stimulus-specific effects (beta values), including the predictors of interest for task and rest conditions, as well as confound predictors derived from fMRIPrep (six motion parameters, motion outliers, physiological parameters from cerebrospinal fluid, white matter and the top five components for anatomical component correction (Behzadi et al., 2007)). The “task vs. rest” contrast computation produced a z-score map, indicating how strongly the BOLD signal differs between the imagery and baseline blocks for each participant and stimulus. Each resulting z-map was first thresholded at the voxel level (p < 0.01, uncorrected) and then corrected for multiple comparisons at the cluster level using 1000 Monte Carlo simulations (cluster-level threshold p < 0.05) to estimate the cluster size threshold. The subsequent second-level analysis was conducted using the SecondLevelModel module in Nilearn (0.9.2). Based on the individual z-score maps from the first-level analysis, a one-sample t-test was applied for each stimulus across all participants. The resulting contrast t-maps were also cluster-level corrected (initial cluster forming threshold p < 0.01, cluster-level threshold p < 0.05).

To explore the sensitivity of different brain regions to multiple emotions, 39 regions which showed more than 50% overlap with the predefined VOI (Supplementary Table S2), based on the automatic anatomical labeling (AAL) atlas (Rolls et al., 2020), were selected for further analysis. The individual z-score map was averaged within different regions of interest (ROIs) and then analyzed across participants. The mean value, SD and distribution were computed across all participants for each ROI and stimulus. Paired t-tests or Wilcoxon signed-rank tests, depending on the distribution normality tested by Shapiro-Wilk tests (p < 0.05), were performed within each ROI between different emotions. The resulting statistics were corrected across all the ROIs using false discovery rate (FDR, p < 0.05).

##### Multivariate pattern analyses

Based on real-time output data, we first examined voxel selection distributions, including selecting frequency across participants and regional selection proportion. The selecting frequency was calculated across individuals after projecting selected voxels onto the standard MNI152 template. The region-level selection proportion was defined as the fraction of selected voxels within each ROI relative to the total selected voxels. These proportion values were then averaged across participants and ranked across regions for visualization.

Pairwise dissimilarities between emotions were then quantified using group-level RDM and its 95% confidence intervals. Derived from the online RDM, dissimilarity values were first converted back to correlation coefficients and transformed using Fisher’s z-transformation. The transformed values were averaged to obtain the mean, and the SD was used to calculate the 95% confidence intervals for z-scores. The mean z-score and its confidence interval bounds were then inversely transformed to obtain Pearson’s r correlation values. Finally, the mean RDM and its confidence intervals were derived by subtracting the corresponding correlation values from 1.

#### 2.5.2 Neurofeedback performance evaluation

Three metrics were defined to evaluate neurofeedback performances, including (1) Pearson’s correlation between the current brain activity pattern and the target base pattern, (2) angular distance between the moving red dot and the target emotion icon on the CSM, and (3) intensity level of the target emotion’s thermometer on the CSM. Pearson’s correlation implicitly measured the actual (dis)similarity between the current and target patterns, while angular distance and intensity level served as direct indicators of (dis)similarity and intensity for the participant. Intensity level exceeded zero only when the current pattern exhibited the highest similarity to the target base pattern.

Group-level evaluation was first conducted over the entire feedback time sequence. The mean and 95% confidence interval of each performance metric across participants were calculated. Specifically for Pearson’s correlation, values relative to all non-target base patterns were also computed. Learning effects within and between runs were examined using linear mixed-effects models (LMMs) implemented in the python function Statsmodels (0.12.2). We excluded the first two feedback time points while fitting the model to avoid low initial values caused by limited data. The model included run, time, and their interaction as fixed effects, with participant as a random intercept. To interpret the significance of the fixed effects within both runs separately, we fit the model twice, each time changing the reference level of the run term. Intercept and linear slope differences between runs were extracted to reflect variations in initial performance and learning rate, respectively. To account for potential non-linearity (e.g., performance plateaus toward the end of a run), additional exploratory models incorporating quadratic or exponential time terms were fit and compared using a likelihood ratio test (LRT).

We also evaluated group-level neurofeedback performance by comparing each metric between the initial and final feedback block (based on the median value across six feedback time points within the block). The statistical significance was tested via paired t-tests or Wilcoxon signed-rank tests, depending on distribution normality tested by Shapiro-Wilk tests (p < 0.05), and then corrected using Bonferroni for two comparisons based on the number of neurofeedback runs.

Individual performance evaluation involved two parts: (1) computing a neurofeedback score for each participant and run, and (2) examining its relationship with participants’ perceived control ratings. Neurofeedback scores were obtained by fitting a linear regression model to each metric’s values over time. Regulation was considered successful for a given metric if the score showed a statistically significant improvement over time, with significance defined as FDR-corrected p < 0.05. Participants’ perceived control ratings, reported on a scale from 0 to 10, reflected their subjective sense of control over brain activity during neurofeedback. To characterize the distribution of the ratings, the mean and SD across participants were calculated for each run, and a paired t-test was conducted to compare ratings between runs. Depending on the distribution normality (Shapiro–Wilk test, p < 0.05), either Pearson’s or Spearman’s correlation was used to assess the relationship between objective neurofeedback scores (i.e., coefficients of the linear regression model) and subjective perceived control ratings.

Furthermore, to visualize individual self-regulation processes, we applied a multidimensional scaling (MDS) (Ciarlo et al., 2022; Goebel et al., 2024; GOWER, 1966; Hefner, 1959; Russo et al., 2021) algorithm to the four base patterns to estimate a two-dimensional representational space for each participant. The real-time neurofeedback patterns were then projected into the same space and displayed as continuous points, reflecting participants’ learning trajectories.

Additionally, we classified participants’ reported emotion imagery strategies into three categories: autobiographical memory recall, imagined situations (i.e., events that did not actually happen or were fictional), and body-related sensations (i.e., awareness of current breathing or bodily feelings).

## 3. RESULTS

All participants (N = 27) completed four localizer runs. Most of them (N = 24) completed two neurofeedback runs targeting joyful relaxation, while one participant completed only one such run. Two participants did not complete any neurofeedback runs due to scanner-related technical issues (N = 1) or personal scheduling conflicts (N = 1). Approximately half (N = 14) of the participants additionally performed an optional neurofeedback run targeting sadness.

### 3.1 Analyses of the localizer data

#### 3.1.1 Univariate patterns of multiple emotions

The results of the group-level univariate analysis for the localizer data are summarized in Figure 2 (see Supplementary Figure S1 for the corresponding analysis for the neurofeedback data). Significant activation clusters, based on the “task vs rest” contrast, were consistently observed across conditions in the prefrontal cortex (PFC; Figure 2A). This activation pattern appears to be consistent with regions typically engaged in emotion imagery and processing.

**Figure 2.**
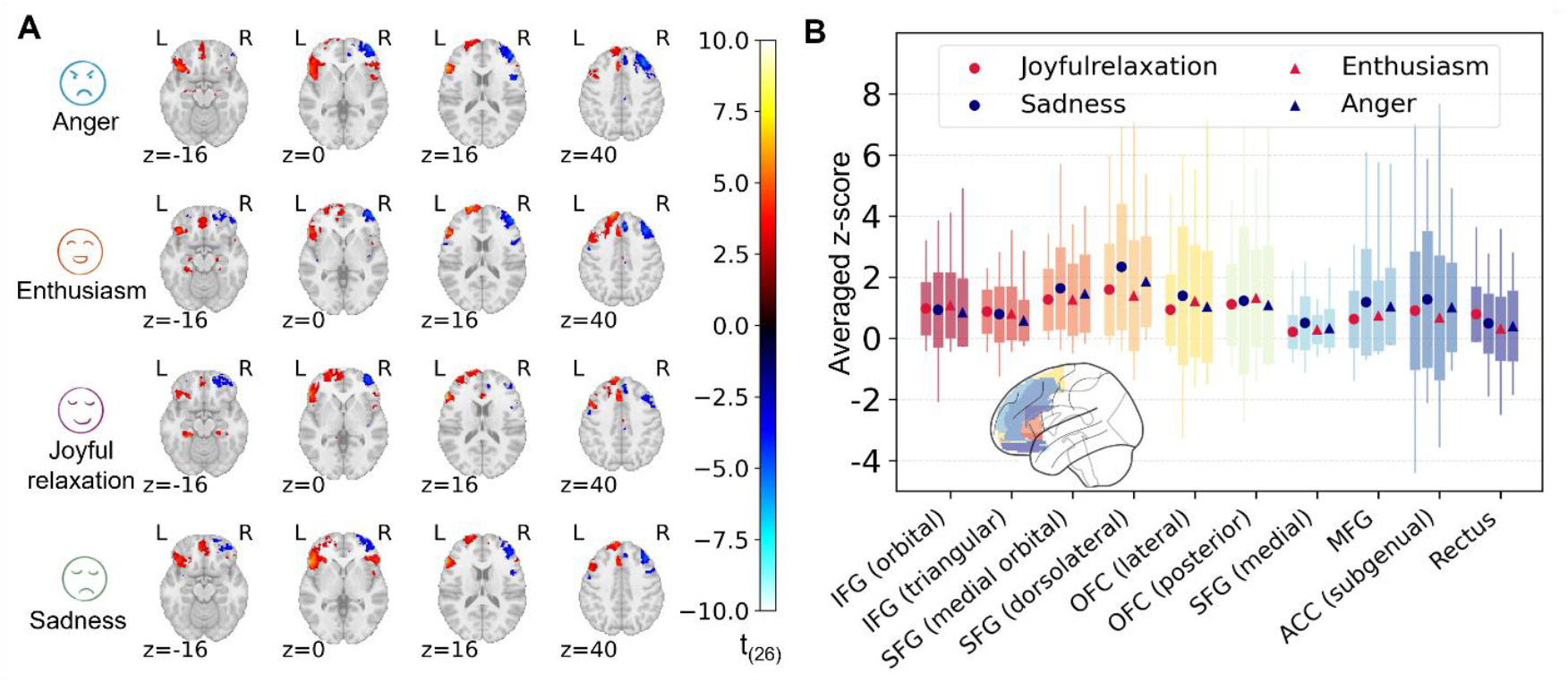
Univariate patterns based on the raw localizer data. (A) T-maps of significant activation for each condition (task vs rest) in MNI space (voxel-level thresholded at p < 0.01, cluster-level corrected at p < 0.05). Only clusters encompassing the target VOI are shown. (B) ROI analysis (left hemisphere), computed by averaging the individual z-maps within each ROI. The boxplot summarizes the mean, SD and distribution of averaged z-scores across participants. The color and shape of each node represent the valence and arousal dimensions of the emotions (circular shape = low arousal, triangular shape = high arousal; red = positive emotions, blue = negative emotions). No region showed significant differences (i.e., p > 0.05) between high and low arousal emotions, or between different positive and negative emotions. MNI = Montreal Neurological Institute; VOI = volume of interest; ROI = region of interest; SD = standard deviation.

In the ROI analysis based on averaged z-values, no significant differences emerged between conditions. This result indicates that simply considering the average activity within each region, as typically assessed in univariate analysis, was insufficient to distinguish between the considered emotions. Results for the 10 ROIs showing the highest group-level activity across conditions in the left hemisphere are illustrated in Figure 2B (see Supplementary Figure S2 for all 39 ROIs). While there was a trend to differentiate positive and negative emotions in dorsolateral superior frontal gyrus (dlSFG) and middle frontal gyrus (MFG), no region showed significant differences. This finding highlights the limitation of univariate analysis in recognizing the neural fingerprints of distinct emotions.

#### 3.1.2 RSA patterns of multiple emotions

The distribution of selected voxels was examined at both voxel and region levels. At the voxel level, the spatial pattern of selected voxels during the online analysis (Figure 3A) closely matched the offline group-level t-maps (Figure 2A). At the ROI level, the regions with the highest selection proportion (Figure 3B) largely overlapped with those showing the strongest activation in the univariate analysis (Figure 2B), primarily involving the frontal gyrus and anterior cingulate cortex (ACC). Based on the base patterns derived from these selected voxels, the group-level averaged RDM distinguished the four target emotions, most prominently along the valence dimension (Figure 3C). A similar pattern could also be observed in the lower and upper bounds of the 95% confidence interval, indicating the robustness of the valence-specific distinctions. Overall, the multivariate results demonstrated the potential of this approach for reliably separating the considered emotional states.

**Figure 3.**
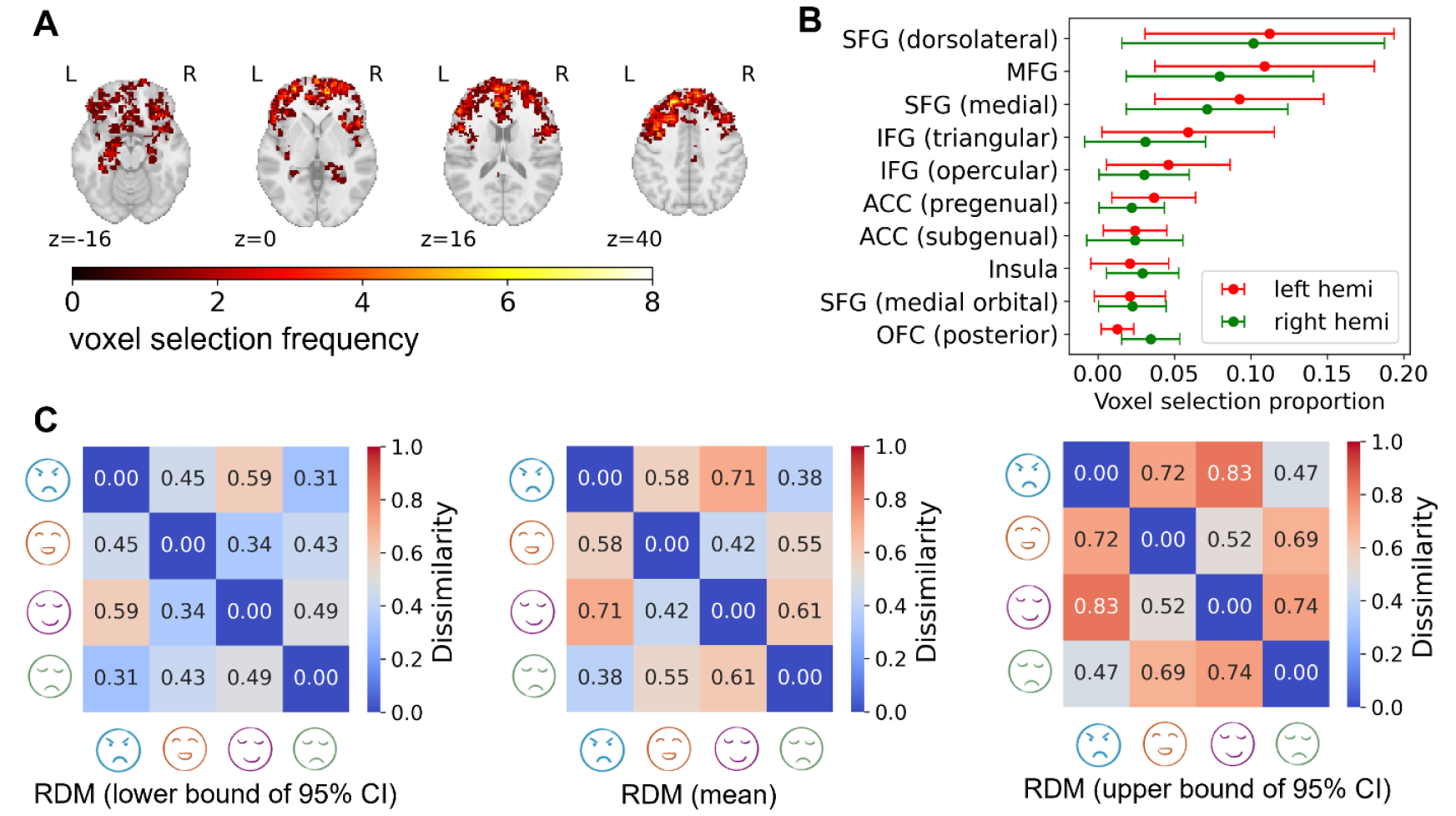
Distribution of selected voxels and RSA patterns based on the real-time localizer data (N = 27). (A) Distribution of the real-time selected voxels. The color shade indicates how often each voxel was selected across participants. (B) Selection probability of each ROI (the top 10) across participants. The x-axis indicates the proportion of selected voxels in each region relative to the total number of selected voxels. (C) Group-level mean RDM and its 95% CIs, derived from the individual online analysis. ROI = region of interest, SFG = superior frontal gyrus, MFG = middle frontal gyrus, IFG = interior frontal gyrus, ACC = anterior cingulate cortex, OFC = orbitofrontal cortex, RDM = representational dissimilarity matrix, CI = confidence interval.

### 3.2 Neurofeedback performance analyses

Here, we report the neurofeedback performance analysis for the emotion of joyful relaxation, as it was completed by the majority of participants in two consecutive runs, allowing for both within-run and between-run performance comparisons. Analysis of the optional third neurofeedback run targeting sadness, which was performed by fewer participants and included a single run, is provided in Supplementary Figure S3-S5 and Table S3-S7.

#### 3.2.1 Group-level neurofeedback performance

##### (1) Full run time-sequence analysis

The neurofeedback performance curves of three metrics over feedback time points are illustrated in Figure 4. In Figure 4A, Pearson’s correlation coefficients between the real-time brain pattern and each of the four emotional base patterns are plotted at every feedback time point, providing a dynamic view of how closely participants’ neural activity aligned with each target state. The correlation with the target base pattern for joyful relaxation became consistently higher than with the other emotional patterns after the first two blocks. This sustained advantage indicates that participants were able to reproducibly engage the target neural pattern, thereby maintaining effective modulation while differentiating it from other emotional states.

**Figure 4.**
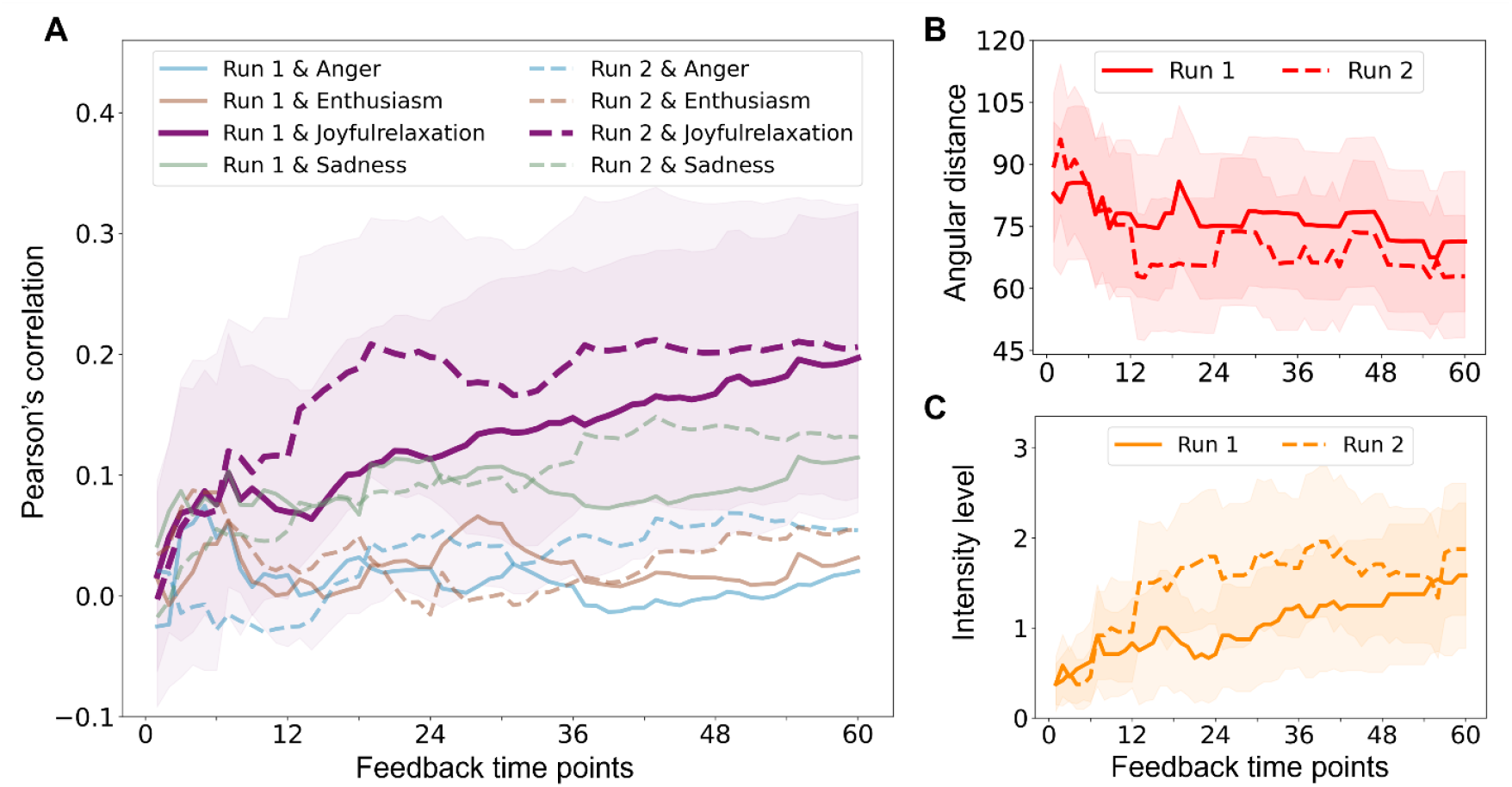
Learning curves of neurofeedback performance metrics (target condition: joyful relaxation). The metrics include Pearson’s correlation (A), angular distance (B), and intensity level (C). In each part, the mean value (across participants) and its 95% confidence interval are plotted over all feedback time points for two runs. Part A additionally shows, in the background, Pearson’s correlations between the real-time brain pattern and each non-target emotional base pattern.

Additionally, as shown in Figure 4A, while correlations with the enthusiasm and anger patterns in the second run showed slight upward trends, all other non-target emotional patterns remained stable over time. In contrast, correlations with the joyful relaxation pattern increased in both runs. The two derivative metrics also showed consistent improvement, reflected by decreasing angular distance (Figure 4B) and increasing intensity level (Figure 4C). The LMM analyses confirmed these trends as statistically significant (Table 1, see Supplementary Table S8 for a complete LMM report), with consistent effects across runs: positive slopes for Pearson’s correlation (Run 1: r = 0.002, p < 0.001; Run 2: r = 0.002, p < 0.001) and intensity level (Run 1: r = 0.017, p < 0.001; Run 2: r = 0.016, p < 0.001), and negative slopes for angular distance (Run 1: r = –0.172, p < 0.001; Run 2: r = –0.232, p < 0.001). These results demonstrate a robust within-run learning effect, reflecting that participants were able to increasingly align their brain activity with the target pattern over time.

**Table 1.** Results of the LMM analysis for both runs (Formula: metric ∼time + Run + time:Run + (1|Participant)). LMM = linear mixed-effects model, R2 = Run 2, R1 = Run1.

|  |  | Pearson's correlation |  |  |  | Angular distance |  |  |  | Intensity level |  |  |  |
| --- | --- | --- | --- | --- | --- | --- | --- | --- | --- | --- | --- | --- | --- |
|  |  | Coef. | Std. Err. | z | p | Coef. | Std. Err. | z | p | Coef. | Std. Err. | z | p |
| **Intercept** | R1 | 0.061 | 0.050 | 1.236 | 0.217 | 81.819 | 7.053 | 11.600 | < 0.001 | 0.525 | 0.293 | 1.794 | 0.073 |
|  | R2 | 0.111 | 0.050 | 2.245 | **0.025** | 77.177 | 7.053 | 10.942 | < 0.001 | 1.025 | 0.293 | 3.499 | < 0.001 |
|  | R2 vs. R1 | 0.050 | 0.013 | 3.710 | < 0.001 | -4.642 | 2.240 | -2.703 | **0.038** | 0.499 | 0.084 | 5.921 | < 0.001 |
| **time** | R1 | 0.002 | 0.000 | 7.432 | < 0.001 | -0.172 | 0.044 | -3.863 | < 0.001 | 0.017 | 0.002 | 9.995 | < 0.001 |
|  | R2 | 0.002 | 0.000 | 6.431 | < 0.001 | -0.232 | 0.044 | -5.221 | < 0.001 | 0.016 | 0.002 | 9.447 | < 0.001 |
|  | R2 vs. R1 | -0.001 | 0.000 | -0.708 | 0.479 | -0.060 | 0.063 | -0.960 | 0.337 | -0.001 | 0.002 | -0.388 | 0.698 |

Figure 4 also revealed distinct patterns in learning dynamics between the two runs. For most time points, both Pearson’s correlation and intensity level were higher in the second run than in the first, while angular distance was consistently lower after the first two blocks. LMM results demonstrated that initial performance in the second run was significantly better across all three metrics, but no significant differences in the linear slopes were observed between runs (Table 1). This pattern suggests that participants seem to begin the second run with an improved baseline regulation ability, likely reflecting carryover from the first run.

Visual inspection of the curves in Figure 4 also suggested a shift in learning patterns: performance in the first run showed a steady progression, while improvements in the second run appeared to plateau after the early blocks. To formally account for these potential non-linear trends, an exploratory model comparison using LRT indicated that including an additional quadratic time term provided a significantly better fit on all three metrics (Pearson’s correlation:

χ^2^(2) = 14.25, p < 0.001; angular distance: χ^2^(2) = 7.10, p = 0.029; intensity level: χ^2^(2) = 70.12, p < 0.001; see Supplementary Table S9). The results from the quadratic model aligned with the observed patterns, revealing significant interactions between run and both the linear (time × run) and quadratic (time^2^ × run) terms across all three metrics (Table 2; Supplementary Table S10 and S11 show complete LMM reports for the quadratic and exponential models, respectively). For Pearson’s correlation and intensity level, the interaction was characterized by a positive linear and negative quadratic term, indicating a sharper initial increase in the second run followed by a deceleration. For angular distance, the interaction showed a negative linear and positive quadratic term, reflecting a steeper initial decline that flattened over time. Overall, these results suggest a potential between-run learning effect, although the exploratory quadratic model adds greater complexity.

**Table 2.**
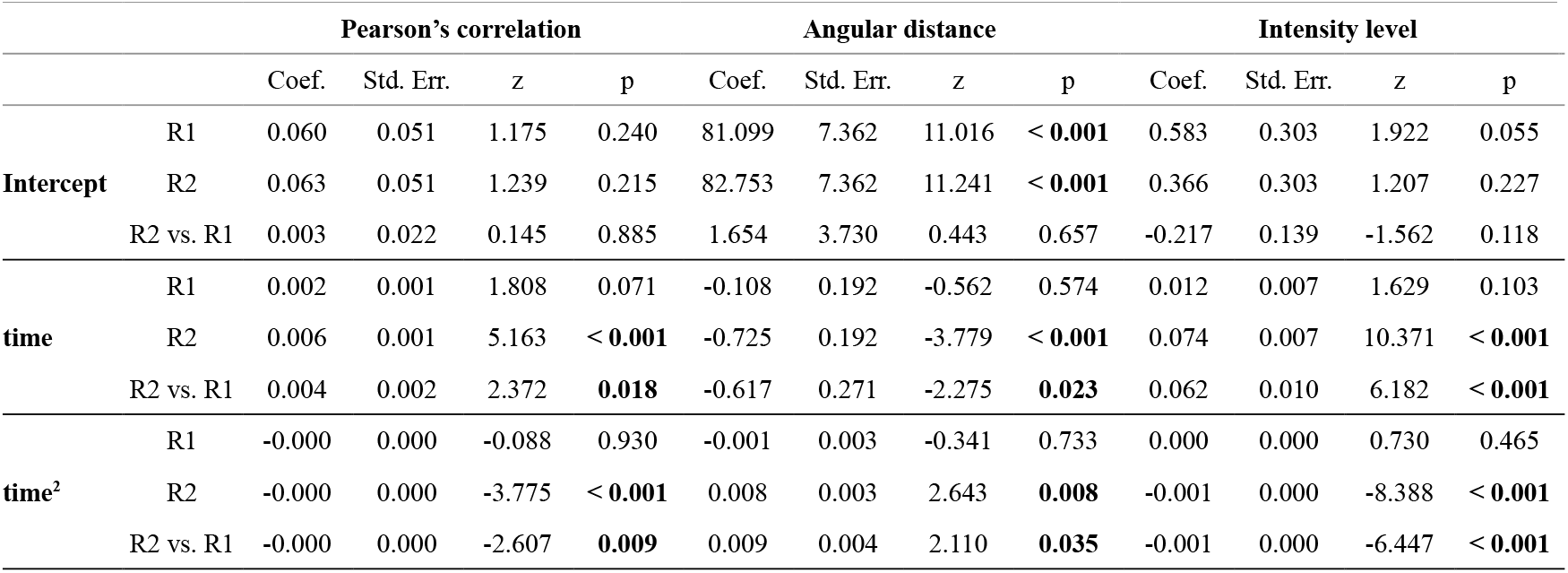
Results of the LMM analysis with quadratic time terms (Formula: metric ∼time + time^2^ + Run + time:Run + time^2^:Run + (1|Participant)). LMM = linear mixed-effects model, R2 = Run 2, R1 = Run1.

|  |  | Pearson's correlation |  |  |  | Angular distance |  |  |  | Intensity level |  |  |  |
| --- | --- | --- | --- | --- | --- | --- | --- | --- | --- | --- | --- | --- | --- |
|  |  | Coef. | Std. Err. | z | p | Coef. | Std. Err. | z | p | Coef. | Std. Err. | z | p |
| **Intercept** | R1 | 0.060 | 0.051 | 1.175 | 0.240 | 81.099 | 7.362 | 11.016 | < **0.001** | 0.583 | 0.303 | 1.922 | 0.055 |
|  | R2 | 0.063 | 0.051 | 1.239 | 0.215 | 82.753 | 7.362 | 11.241 | < **0.001** | 0.366 | 0.303 | 1.207 | 0.227 |
|  | R2 vs. R1 | 0.003 | 0.022 | 0.145 | 0.885 | 1.654 | 3.730 | 0.443 | 0.657 | -0.217 | 0.139 | -1.562 | 0.118 |
| **time** | R1 | 0.002 | 0.001 | 1.808 | 0.071 | -0.108 | 0.192 | -0.562 | 0.574 | 0.012 | 0.007 | 1.629 | 0.103 |
|  | R2 | 0.006 | 0.001 | 5.163 | < **0.001** | -0.725 | 0.192 | -3.779 | < **0.001** | 0.074 | 0.007 | 10.371 | < **0.001** |
|  | R2 vs. R1 | 0.004 | 0.002 | 2.372 | **0.018** | -0.617 | 0.271 | -2.275 | **0.023** | 0.062 | 0.010 | 6.182 | < **0.001** |
| **time<sup>2</sup>** | R1 | -0.000 | 0.000 | -0.088 | 0.930 | -0.001 | 0.003 | -0.341 | 0.733 | 0.000 | 0.000 | 0.730 | 0.465 |
|  | R2 | -0.000 | 0.000 | -3.775 | < **0.001** | 0.008 | 0.003 | 2.643 | **0.008** | -0.001 | 0.000 | -8.388 | < **0.001** |
|  | R2 vs. R1 | -0.000 | 0.000 | -2.607 | **0.009** | 0.009 | 0.004 | 2.110 | **0.035** | -0.001 | 0.000 | -6.447 | < **0.001** |

##### (2) Initial versus final NF block comparison

Within each neurofeedback run, comparisons between the initial and final blocks revealed no significant changes in Pearson’s correlation with the target base pattern (Run 1: t_(23)_ = 1.811, p = 0.166; Run 2: t_(23)_ = 1.911, p = 0.137; Figure 5A) or in angular distance to the target condition (Run 1: W = 119.0, p = 0.780; Run 2: W = 81.0, p = 0.098; Figure 5B). On the other hand, intensity level started near zero and ended significantly higher than the initial level (Run 1: W = 10.5, p = 0.028; Run 2: W = 7.0, p = 0.005; Figure 5C). This initial-final within-run comparison indicated a trend toward improved performance from the beginning to the end of each run, though not all metrics reached statistical significance. The lack of significance in some metrics may be related to substantial inter-individual variability, as reflected by the relatively large SDs across participants in both runs (mean ± SD for initial vs. final; Pearson’s correlation: 0.06 ± 0.22 vs. 0.18 ± 0.29 in Run 1, 0.05 ± 0.28 vs. 0.19 ± 0.29 in Run 2; angular distance: 85.47 ± 45.40 vs. 71.24 ± 42.33 in Run 1, 91.30 ± 44.46 vs. 62.78 ± 37.14 in Run 2; intensity level: 0.48 ± 0.82 vs. 1.50 ± 2.02 in Run 1; 0.42 ± 0.69 vs. 1.85 ± 1.82 in Run 2; Figure 5).

**Figure 5.**
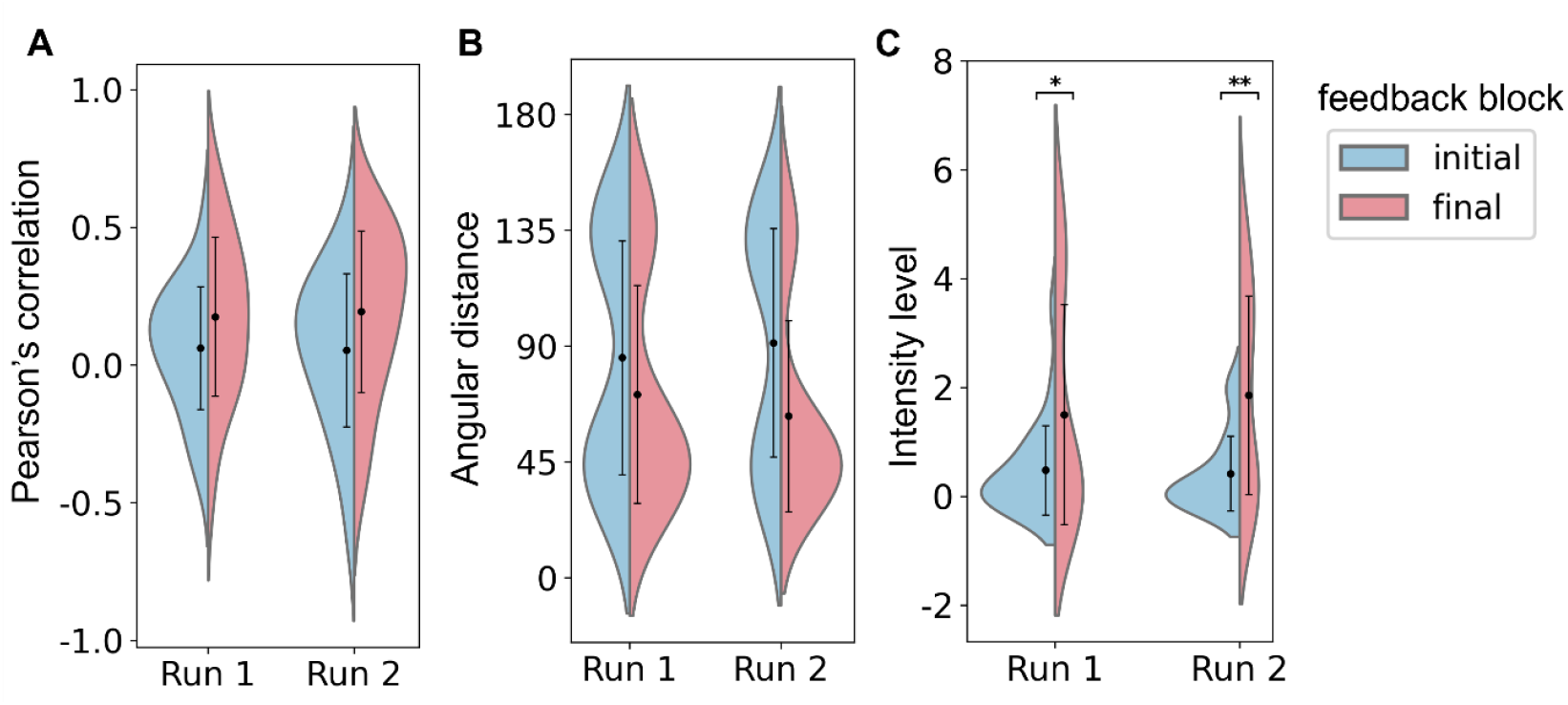
Comparison of neurofeedback performance metrics between initial and final feedback blocks (target condition: joyful relaxation). The median value of each metric within the corresponding block was extracted and compared. (A) Pearson’s correlation coefficient between target pattern and the pattern extracted in the initial and final feedback blocks. (B) Angular distance in the initial and final feedback block, measured by calculating the angular distance between the red dot and the target stimulus on the CSM. (C) Same as (B) but using the thermometer reading of the target stimulus on the CSM as metric of intensity level. The violin plot shows the distribution of each metric across participants, and the error bar shows the corresponding mean and SD. Paired t-tests or Wilcoxon signed-rank tests were performed on the initial and final metrics. Results were considered significant (*: p < 0.05, **: p < 0.01) after Bonferroni correction (n = 2). CSM = circular semantic map; SD = standard deviation.

#### 3.2.2 Subjectively perceived control and neurofeedback performance

At the end of each neurofeedback run, participants rated their perceived control over brain activity on a scale from 0 to 10 and reported the imagery strategies they had used, providing a subjective reference for their neurofeedback performance. Three sets of ratings were missing, two of which were from participants who had completed both joyful relaxation runs.

Perceived control ratings demonstrated similar distributions across runs, with descriptive statistics (mean ± SD) of 4.64 ± 1.62 for the first run and 5.45 ± 2.28 for the second run. A paired t-test revealed no significant difference between runs (t_(21)_ = 1.80, p = 0.086), although there was a trend toward higher ratings in the second run. Our offline individual neurofeedback performance analyses showed that 22 out of 24 participants achieved successful regulation on at least one metric and 10 of them succeeded on all three metrics (see Supplementary Table S12), objectively reflecting their ability to control mental states.

Furthermore, as shown in Figure 6, correlation analyses revealed a significant positive relationship between participants’ perceived control ratings and their performance slopes for Pearson’s correlation. In contrast, no significant relationships were observed for angular distance or intensity level. These findings were consistent across both neurofeedback runs, supporting the alignment between subjective reports and objective performance measures.

**Figure 6.**
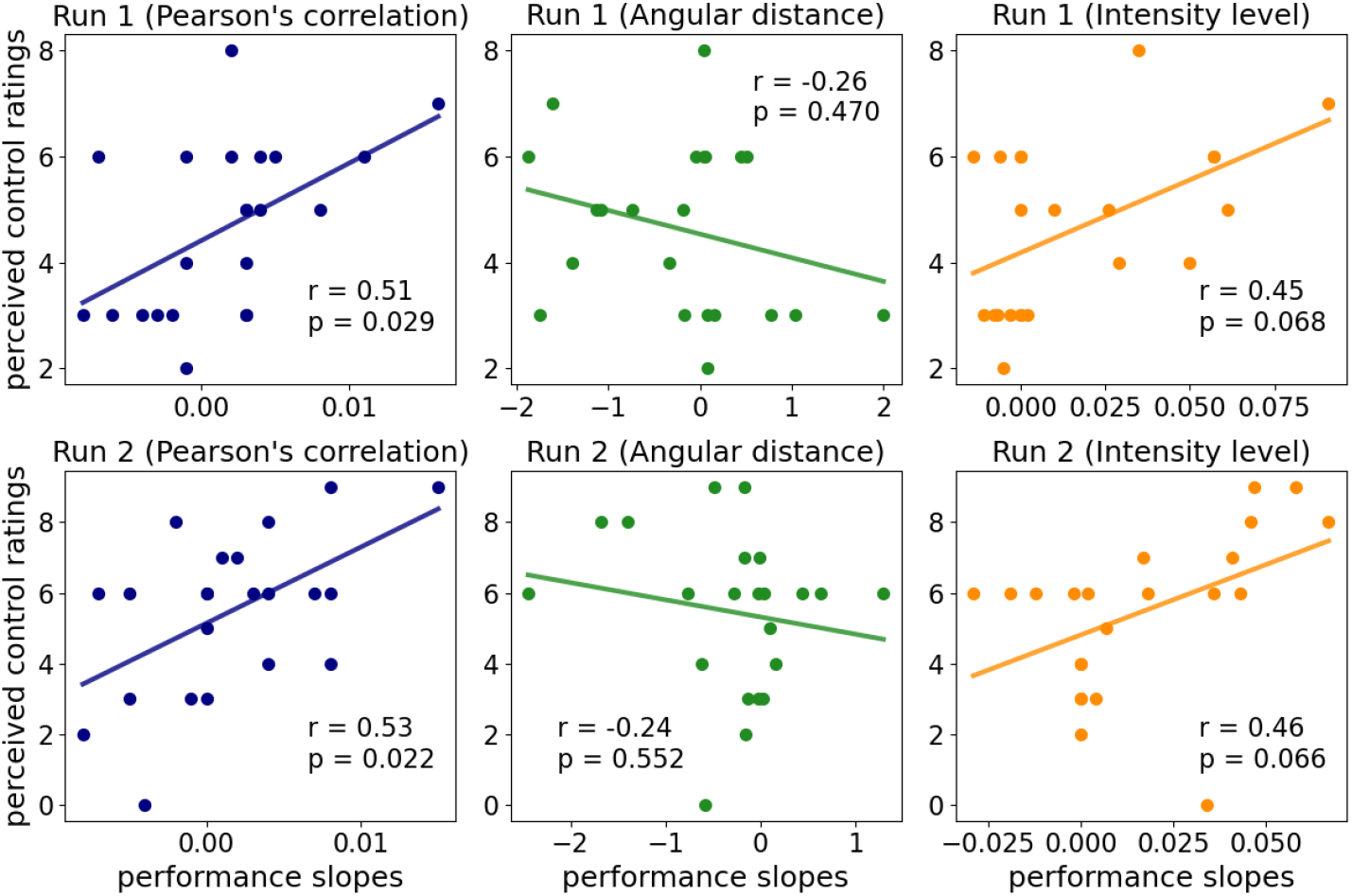
Correlation analyses between participants’ perceived control ratings and neurofeedback performance slopes for the condition joyful relaxation (N = 22). The perceived control ratings (y-axis) reflect participants’ perceived control over brain activity, rated on a scale from 0 to 10. The performance slopes (x-axis) were obtained from individual linear regression models across the feedback time sequence. Statistical values were calculated using either Pearson’s or Spearman’s correlation, with Bonferroni correction across runs (n = 2) applied.

In addition, the MDS-based representational space was used to visualize descriptively each participant’s modulation trajectory during neurofeedback runs (Supplementary Figure S6, S7). Although some trajectories diverged, most aligned with participants’ perceived control ratings. Specifically, higher ratings were often accompanied by neurofeedback sequences that move progressively toward the corresponding base emotion as determined by the localizer. The distribution of emotion imagery strategies across participants is provided in Supplementary Figure S8, with most participants relying on autobiographical recall to induce the target emotional states.

## DISCUSSION

Here we investigated for the first time the feasibility of RSA-based rt-fMRI-sNF to navigate toward a predefined emotional state. To overcome the limitations of conventional univariate rt-fMRI-NF studies, we computed both the similarity and intensity of brain activity pattern using RSA as multivariate analysis method and delivered them to participants by a novel feedback visualization approach, the CSM. Our study provided four main insights: First, data from the localizer showed that RSA was able to differentiate between four different emotional states induced by mental imagery, providing for the first time proof-of-concept for the statistical sensitivity of RSA when used with the parameter choices made in our study. Second, participants were able to navigate their mental state toward a target state during consecutive neurofeedback training, providing evidence for the feasibility of our approach. Third, an analysis of participants’ neurofeedback performance (based on the outcome metrics of correlation, similarity and intensity) from two neurofeedback runs suggested that self-regulation performance improved over time, indicating that participants learned to improve controlling the CSM-based visual feedback. Lastly, data showed that overall participants reported moderate control over the feedback, and this perceived control ratings were significantly correlated with the correlation metric, consistently across both neurofeedback runs.

Our pipeline included both univariate and multivariate processing steps, allowing us to compare results for both types of analyses and their performance in distinguishing between multiple affective states. In conventional univariate rt-fMRI-NF studies (Paret et al., 2019; Weiskopf et al., 2004), participants receive feedback based on the average activity within a predefined target region and learn to modulate this activity to influence the feedback intensity. While effective in simpler paradigms (Barreiros et al., 2019; Johnston et al., 2011), this approach becomes limited when applied to studies involving multiple emotional states (Habes et al., 2013), as univariate methods have difficulty in capturing emotion-specific neural patterns, since different emotions often activate overlapping brain regions (Shibata et al., 2016; Todd et al., 2020). We also observed this limitation in our own data when performing univariate ROI analysis, such that the distributions of individual averaged contrast scores (task vs rest) showed substantial overlap and no significant differences across multiple emotional states for the selected emotion-related ROIs (see Supplementary Figure S2). In contrast, multivariate methods analyze distributed multi-voxel brain activity patterns (Haxby, 2012; Mahmoudi et al., 2012), allowing for capturing the complex neural differences involved in processing multiple emotions. In our findings, the voxels contributing most to these patterns were primarily located in the dorsomedial and dorsolateral PFC (dmPFC, dlPFC), ACC, insula, and orbitofrontal cortex (OFC), which were found to be consistently implicated in emotion imagery in previous studies (Linhartová et al., 2019). Furthermore, the group-level averaged RDM revealed clear distinctions between positive and negative emotional states. These findings show that our real-time RSA-based approach can reliably distinguish between multiple emotions, making it better suited than univariate methods for neurofeedback studies targeting complex emotional states.

Importantly, we extended the rt-fMRI-sNF paradigm, previously tested using mental imagery of objects (Ciarlo et al., 2022; Russo et al., 2021), to the domain of emotion regulation. Object imagery involves the visualization of concrete external items (e.g., cats and chairs), relying on relatively objective and consistent visual-spatial representations (Sima et al., 2013). In contrast, emotion imagery requires the mental simulation of internal emotional experiences, which are inherently subjective and influenced by personal context and memory (Wicken et al., 2021). The added cognitive and affective complexity makes emotion imagery more challenging to engage and regulate. Our group-level offline assessment of neurofeedback performance supports the feasibility of applying this paradigm to emotion regulation. Specifically, we observed a reduced angular distance between the real-time activity pattern and the target emotion pattern, as well as increased activation intensity in the final emotion imagery block compared to the first. Besides, within-run learning trends showed significant and consistent improvement across three evaluation metrics. These results suggest that participants were able to progressively align their brain activity with the target emotional states, demonstrating effective engagement with visual feedback. Moreover, we observed higher initial performance (from linear model) and larger learning rate (from exploratory quadratic model) in the second neurofeedback run. This suggests that participants may have retained effective strategies from the first run, enabling more efficient engagement during the subsequent run.

In addition, we examined the effectiveness of the CSM-based visual feedback, a novel feedback presentation method, in the context of emotion regulation training. Conventional univariate rt-fMRI-NF paradigms typically use a thermometer-like display to reflect brain activation intensity. The previous rt-fMRI-sNF paradigm for object imagery (Ciarlo et al., 2022; Russo et al., 2021) focused pattern similarity by projecting the real-time and target activity patterns into a two-dimensional space according to the pairwise dissimilarities. Building on this approach and its variant (Goebel et al., 2024), the CSM placed multiple thermometers and a moving red dot in a circular arrangement, providing feedback on both pattern similarity and activation intensity. Our individual neurofeedback performance results indicated successful learning trends in most participants (see Supplementary Table S12). Of the 24 participants, 22 achieved successful regulations in at least one metric during one or more neurofeedback runs, and 10 participants succeeded across all three metrics during at least one neurofeedback run. These findings suggest that participants were able to engage with and benefit from the feedback, supporting the practical usability of the CSM-based visual feedback for individual-level emotion regulation. This notion is further supported by the relationship between participants’ subjective perceptions and their objectively measured ability to regulate brain activity through neurofeedback. Participants generally reported moderate perceived control, suggesting a consistent subjective experience during the experiment. While perceived control ratings showed no significant difference between runs, exploratory descriptive analyses using MDS indicated that for most participants with moderate to high control ratings (>6), the similarity between neurofeedback and base patterns increased over time (see Supplementary Figure S6, S7). Importantly, we observed a significant positive association between perceived control ratings and neurofeedback performance on the Pearson’s correlation metric. This alignment between subjective perception and objectively calculated performance further underscores the feasibility of the rt-fMRI-sNF paradigm, indicating that participants can meaningfully interpret and utilize feedback to modulate neural activity.

Furthermore, we investigated group-level activation patterns during the localizer and neurofeedback runs (see Supplementary Figure S1). The localizer task, focusing on emotion generation, primarily engaged the prefrontal regions, including the dlPFC, dmPFC and OFC. On the other hand, the neurofeedback runs, emphasizing emotion regulation, showed descriptively stronger activation in the insula and ventromedial PFC (vmPFC). These descriptive differences align with prior research on the neural correlates of emotion generation and regulation. Specifically, the vmPFC is extensively involved in modulating affective responses and integrating feedback to guide emotional adjustments (Alexander et al., 2023). The insula, especially its anterior portion, plays a key role in interoceptive awareness, enabling individuals to monitor and regulate internal emotional states based on real-time feedback (Menon & Uddin, 2010).

While the presented data suggests that our approach has overcome several limitations of traditional univariate neurofeedback studies in the context of emotion regulation, several aspects remain to be addressed in future work.

One key limitation of this study is the insufficient data for neurofeedback performance comparisons across emotions. While 24 participants completed repeated neurofeedback runs for the emotion of joyful relaxation, only 14 participants performed a single run for sadness, limiting direct comparisons of neurofeedback performance between emotional conditions. The rt-fMRI-sNF paradigm requires multiple functional localizer runs to generate base patterns prior to neurofeedback training, which limits the number of neurofeedback runs that can be performed within a single session. Besides, a long measurement may lead to participant fatigue and reduce the mental imagery quality in emotion-based studies. For instance, when participants become tired or uncomfortable, it may be particularly challenging for them to generate and regulate high-arousal positive emotions effectively. A potential solution to this constraint is adopting a multi-session design, in which localizer and neurofeedback runs are performed on separate days to allow for more extensive training and data collection. Depending on the study design, such an approach would likely require precise alignment of functional localizer data across measurement sessions to ensure comparability (Frost & Goebel, 2013; Haxby et al., 2020). Moreover, multi-session neurofeedback training protocols will have to deal with potential challenges related to the test–retest reliability of emotion processing (Berboth et al., 2021; Nord et al., 2017) and, in particular, emotional mental imagery. However, a multivariate approach likely also entails an improvement in relation to this challenge compared to multi-session univariate emotion regulation neurofeedback paradigms (Noble et al., 2021).

As a hardware-related limitation, we used a voxel size of 3mm^3^ at 3 Tesla combined with spatial smoothing of the functional data (6mm Gaussian kernel). The resulting effective resolution might not optimally support differentiating overlapping patterns of different emotions. It remains to be seen whether the increased signal to noise and/or higher spatial resolution of 7 Tesla ultra-high field fMRI will result in substantial improvements in rt-RSA neurofeedback. While not as widespread as 3 Tesla scanners, clinical 7 Tesla devices are becoming increasingly available, which could be relevant for future clinical applications of semantic neurofeedback of emotions.

Lastly, between-subject variability in emotional mental imagery and its influence on neurofeedback performance warrant further investigation. To provide a personalized training approach, our employed real-time RSA-based method selected individualized target voxels and generated participant-specific RDMs. However, we observed varying output RDMs (Figure 3C) across participants during the localizer, which may have introduced differences in regulation difficulty among participants. Incorporating a screening step to assess baseline mental imagery capabilities, along with enhanced pre-experimental training, may help reduce the effects of individual differences and yield improved neurofeedback performance.

## CONCLUSION

This study explored the feasibility of an rt-fMRI-sNF paradigm for guiding emotion regulation. By incorporating an RSA-informed CSM to visualize both the similarity and intensity of brain activity in real time, participants were able to navigate among distinct emotional states using mental imagery. Our localizer findings showed the effectiveness of RSA in differentiating overlapping emotional representations, highlighting its potential to capture the complexity of affective brain patterns. Group-level analyses revealed performance improvements within each neurofeedback run and higher initial performance in the second run, suggesting that participants can effectively modulate and maintain target emotional states when guided by RSA-informed feedback. At the individual level, most participants achieved successful regulation on at least one performance metric, and their perceived control ratings were positively associated with objective neurofeedback outcomes. Overall, this paradigm offers a promising multivariate framework for investigating and training emotion regulation through neurofeedback.

## Supporting information

CRED-nf checklist

## Acknowledgements

This research was funded by the RWTH Aachen University Exploratory Research Space (ERS) OPEN Seed Fund (OPFS727), the START Program of the Faculty of Medicine, RWTH Aachen, and the Junior Principal Investigator (JPI) fellowship funded by the Excellence Strategy of the Federal Government and the Laender (Grant No. JPI074-21). Xuelei Wang was funded by the China Scholarship Council (No. 202206060018). The study was also supported by the Brain Imaging Facility of the Interdisciplinary Center for Clinical Research (IZKF) within the Faculty of Medicine at RWTH Aachen University.

## Data Availability Statement

The data supporting the findings of this study will be made publicly available in a suitable repository at a later date.

