## Supplementary material for "Toward navigating emotional states using real-time representational similarity analysis fMRI neurofeedback - a feasibility study": CRED-nf checklist

### CRED-nf checklist summary

*22 September, 2025*

**Manuscript title:** Toward navigating emotional states using real-time representational similarity analysis fMRI neurofeedback - a feasibility study
**Corresponding Author:** David M. A. Mehler; Jana Zweerings

| Item No. | Checklist item | Manuscript Details |
|---|---|---|
| **Pre-experiment** | | |
| 1a | Pre-register experimental protocol and planned analyses | This feasibility study was preregistered (https://osf.io/5ybuw). |
| 1b | Justify sample size | Because this work was designed as a feasibility study, no formal a priori power analysis was conducted; instead, sample size was determined based on practical constraints and the goal of evaluating methodological feasibility. |
| **Control groups** | | |
| 2a | Employ control group(s) or control condition(s) | *This experiment did not include a control group or control condition* |
| 2b | When leveraging experimental designs where a double-blind is possible, use a double-blind | *NA: A double-blind was not appropriate for this experiment* |
| 2c | Blind those who rate the outcomes | *NA: There was only one participant group* |
| | Blind those who analyse the data | *NA: There was only one participant group* |
| 2d | Examine to what extent participants and experimenters remain blinded | *NA: There was only one participant group* |
| 2e | In clinical efficacy studies, employ a standard-of-care intervention group as a benchmark for improvement | *NA: This is not a clinical efficacy study* |
| **Control measures** | | |
| 3a | Collect data on psychosocial factors | Prior to and after the measurement, all participants completed questionnaires assessing their current mood state and their ability to regulate emotions, as measured by the Profile Of Mood States (POMS) (McNair, 1971) and Emotion Regulation Questionnaire (ERQ) (Gross & John, 2003). |

*This report has been generated using the CRED-nf online checklist version 1.0, adapted from the CRED-nf manuscript which is freely available here https://doi.org/10.1093/brain/awaa009.*

| | | |
|---|---|---|
| 3b | Report whether participants were provided with a strategy | They were then asked to use mental imagery to separately induce and intensify each of the four emotional states based on their own subjective experience, without receiving any specific imagery instructions beforehand. |
| 3c | Report the strategies participants used | Additionally, we classified participants' reported emotion imagery strategies into three categories: autobiographical memory recall, imagined situations (i.e., events that did not actually happen or were fictional), and body-related sensations (i.e., awareness of current breathing or bodily feelings). The distribution of emotion imagery strategies across participants is provided in Supplementary Figure S7. |
| 3d | Report methods used for online-data processing and artifact correction | Data was processed online using Turbo-BrainVoyager (TBV) version 4.2 (Brain Innovation B.V., Maastricht, The Netherlands) and imported in Python using the TBV Network Access Plugin and the corresponding interface (https://github.com/expyriment/expyriment-stash/) as part of the open-source Expyriment library. The rt-RSA Python tool, freely available at (https://github.com/assuntaciarlo/rtRSA_GUI), was used for pattern analysis and feedback display. |
| 3e | Report condition and group effects for artifacts | *Condition and group effects for artifacts were not measured, or not reported in the manuscript* |
| **Feedback specifications** | | |
| 4a | Report how the online-feature extraction was defined | After all the four localizer runs, the rt-RSA tool was used to generate four base patterns by selecting voxels which were related to emotion imagination (univariate selection) and allowed semantic discernibility between emotions (multivariate selection). In the univariate selection, voxels were picked based on their absolute t-values, specifically those whose absolute t-values ranked within the top 60% to 90% of all the voxels within the defined emotion network. The multivariate selection first used a searchlight approach (with a spherical radius of 1 voxel) to generate a four-by-four representational dissimilarity matrix (RDM) for each voxel of the initial mask obtained from the univariate selection. This RDM encoded the dissimilarities between the four emotional patterns obtained from each searchlight, defined as one minus Pearson's correlation coefficient. Voxels were then selected if their RDMs achieved the top 5% similarity to a predefined model RDM, which treated emotions of opposing valence as maximally dissimilar while disregarding differences in arousal. |
| 4b | Report and justify the reinforcement schedule | *The manuscript does not report or justify the reinforcement schedule* |

| | | |
|---|---|---|
| 5b | Plot within-session and between-session regulation blocks of feedback variable(s), as well as pre-to-post resting baselines or contrasts | Additionally, as shown in Figure 4A, while correlations with the enthusiasm and anger patterns in the second run showed slight upward trends, all other non-target emotional patterns remained stable over time. In contrast, correlations with the joyful relaxation pattern increased in both runs. These results demonstrate a robust within-run learning effect, reflecting that participants were able to increasingly align their brain activity with the target pattern over time. This pattern suggests that participants seem to begin the second run with an improved baseline regulation ability, likely reflecting carryover from the first run. Overall, these results suggest a potential between-run learning effect, although the exploratory quadratic model adds greater complexity. |
| 5c | Statistically compare the experimental condition/group to the control condition(s)/group(s) (not only each group to baseline measures) | *The manuscript does not statistically compare the experimental condition/group to the control condition(s)/group(s)* |
| **Outcome measures - behaviour** | | |
| 6a | Include measures of clinical or behavioural significance, defined a priori, and describe whether they were reached | Perceived control ratings demonstrated similar distributions across runs, with descriptive statistics (mean $\pm$ SD) of 4.64 $\pm$ 1.62 for the first run and 5.45 $\pm$ 2.28 for the second run. A paired t-test revealed no significant difference between runs ($t(21) = 1.80$, $p = 0.086$), although there was a trend toward higher ratings in the second run. |
| 6b | Run correlational analyses between regulation success and behavioural outcomes | Furthermore, as shown in Figure 6, correlation analyses revealed a significant positive relationship between participants' perceived control ratings and their performance slopes for Pearson's correlation. In contrast, no significant relationships were observed for angular distance or intensity level. These findings were consistent across both neurofeedback runs, supporting the alignment between subjective reports and objective performance measures. |
| 7a | Upload all materials, analysis scripts, code, and raw data used for analyses, as well as final values, to an open access data repository, when feasible | The rt-RSA Python tool, freely available at (https://github.com/assuntaciarlo/rtRSA_GUI), was used for pattern analysis and feedback display. The offline analysis scripts can be found at (https://github.com/xuelei-1021/rsanf). The data supporting the findings of this study will be made publicly available in a suitable repository at a later date. |

*This report has been generated using the CRED-nf online checklist version 1.0, adapted from the CRED-nf manuscript which is freely available here https://doi.org/10.1093/brain/awaa009.*
