## supplementary information for "Toward navigating emotional states using real-time representational similarity analysis fMRI neurofeedback - a feasibility study"

### Supplementary Materials

**Table S1.** Predefined emotion-related regions. Subcortical regions, including amygdala and hippocampus, are based on the Harvard-Oxford atlas (maximum probability maps, thresholded at 25%, 1 mm resolution). Cortical regions are based on the Schaefer atlas (17-network parcellation, 200 parcels, 1 mm resolution). The coordinates shown as '-' indicate that the corresponding labels were not defined in that hemisphere.

| Label Name | Network Name | Region Name | MNI Coordinates | |
|---|---|---|---|---|
| | | | Left Hemisphere | Right Hemisphere |
| Amygdala | - | amygdala | (-23, -5, -18) | (23, -4, -18) |
| Hippocampus | - | hippocampus | (-25, -22, -14) | (26, -21, -14) |
| ContA_Cingm_1 | control A | middle cingulate cortex | (-3, 4, 30) | (5, 3, 30) |
| ContA_PFCd_1 | control A | dorsal prefrontal cortex | (-22, 6, 62) | (26, 7, 58) |
| ContA_PFCl_1 | control A | lateral prefrontal cortex | (-44, 20, 27) | (52, 11, 21) |
| ContA_PFCl_2 | control A | lateral prefrontal cortex | (-48, 6, 29) | (46, 24, 26) |
| ContA_PFCl_3 | control A | lateral prefrontal cortex | (-43, 6, 43) | - |
| ContA_PFClv_1 | control A | lateral ventral prefrontal cortex | (-42, 40, 16) | - |
| ContB_PFCld_1 | control B | lateral dorsal prefrontal cortex | - | (41, 33, 37) |
| ContB_PFCld_2 | control B | lateral dorsal prefrontal cortex | - | (42, 14, 49) |
| ContB_PFCld_3 | control B | lateral dorsal prefrontal cortex | - | (23, 24, 53) |
| ContB_PFCl_1 | control B | lateral prefrontal cortex | (-40, 19, 49) | - |
| ContB_PFClv_1 | control B | lateral ventral prefrontal cortex | (-42, 49, -6) | (36, 46, -13) |
| ContB_PFClv_2 | control B | lateral ventral prefrontal cortex | (-28, 58, 8) | (29, 58, 5) |
| ContB_PFCmp_1 | control B | medial posterior prefrontal cortex | - | (7, 25, 55) |
| ContC_Cingp_1 | control C | posterior cingulate cortex | (-5, -29, 28) | (5, -24, 31) |
| DefaultA_PFCd_1 | default A | dorsal prefrontal cortex | (-24, 25, 49) | (29, 30, 42) |
| DefaultA_PFCm_1 | default A | medial prefrontal cortex | (-6, 36, -10) | (8, 42, 4) |
| DefaultA_PFCm_2 | default A | medial prefrontal cortex | (-12, 63, -6) | (6, 29, 15) |
| DefaultA_PFCm_3 | default A | medial prefrontal cortex | (-6, 44, 7) | (8, 58, 18) |
| DefaultB_PFCd_1 | default B | dorsal prefrontal cortex | (-8, 59, 21) | (15, 46, 44) |
| DefaultB_PFCd_2 | default B | dorsal prefrontal cortex | (-11, 47, 45) | - |
| DefaultB_PFCd_3 | default B | dorsal prefrontal cortex | (-3, 33, 43) | - |
| DefaultB_PFCd_4 | default B | dorsal prefrontal cortex | (-9, 17, 63) | - |
| DefaultB_PFCv_1 | default B | ventral prefrontal cortex | (-35, 20, -13) | (51, 28, 0) |
| DefaultB_PFCv_2 | default B | ventral prefrontal cortex | (-32, 42, -13) | - |
| DefaultB_PFCv_3 | default B | ventral prefrontal cortex | (-46, 31, -7) | - |
| DefaultB_PFCv_4 | default B | ventral prefrontal cortex | (-52, 22, 8) | - |
| DefaultC_PHC_1 | default C | parahippocampal cortex | (-26, -32, -18) | (28, -36, -14) |
| LimbicB_OFC_1 | limbic B | orbital frontal cortex | (-24, 22, -20) | (12, 39, -22) |
| LimbicB_OFC_2 | limbic B | orbital frontal cortex | (-10, 35, -21) | (28, 22, -19) |
| LimbicB_OFC_3 | limbic B | orbital frontal cortex | - | (5, 37, -14) |
| LimbicB_OFC_4 | limbic B | orbital frontal cortex | - | (15, 64, -8) |
| SalVentAttnA_FrOper_1 | salience / ventral attention A | frontal operculum | (-39, 1, 11) | (43, 7, 4) |
| SalVentAttnA_FrOper_2 | salience / ventral attention A | frontal operculum | (-51, 9, 11) | - |

| Label Name | Network Name | Region Name | MNI Coordinates | |
|---|---|---|---|---|
| | | | Left Hemisphere | Right Hemisphere |
| SalVentAttnA_Ins_1 | salience / ventral attention A | insula | (-39, -4, -4) | (41, 6, -15) |
| SalVentAttnA_Ins_2 | salience / ventral attention A | insula | - | (46, -4, -4) |
| SalVentAttnB_Ins_1 | salience / ventral attention B | insula | (-33, 20, 5) | (34, 21, -8) |
| SalVentAttnB_Ins_2 | salience / ventral attention B | insula | - | (36, 24, 5) |
| SalVentAttnB_PFCl_1 | salience / ventral attention B | lateral prefrontal cortex | (-28, 43, 31) | (30, 48, 27) |
| SalVentAttnB_PFClv_1 | salience / ventral attention B | lateral ventral prefrontal cortex | - | (43, 45, 10) |
| SalVentAttnB_PFCmp_1 | salience / ventral attention B | medial posterior prefrontal cortex | (-6, 30, 25) | (7, 31, 28) |

**Table S2.** Regions used for offline analysis (more than 50% overlapped with the predefined regions, based on AAL3 atlas). AAL = automatic anatomical labeling.

| Anatomical description | AAL3 label | overlap/% | AAL3 label | overlap/% |
|---|---|---|---|---|
| Superior frontal gyrus, dorsolateral | Frontal_Sup_2_L | 70.76 | Frontal_Sup_2_R | 65.00 |
| Middle frontal gyrus | Frontal_Mid_2_L | 74.42 | Frontal_Mid_2_R | 74.88 |
| Inferior frontal gyrus, opercular part | Frontal_Inf_Oper_L | 79.38 | Frontal_Inf_Oper_R | 78.34 |
| Inferior frontal gyrus, triangular part | Frontal_Inf_Tri_L | 81.02 | Frontal_Inf_Tri_R | 68.99 |
| Inferior frontal gyrus pars orbitalis | Frontal_Inf_Orb_2_L | 83.91 | Frontal_Inf_Orb_2_R | 69.68 |
| Olfactory cortex | Olfactory_L | 71.48 | Olfactory_R | 77.78 |
| Superior frontal gyrus, medial | Frontal_Sup_Medial_L | 72.03 | Frontal_Sup_Medial_R | 78.12 |
| Superior frontal gyrus, medial orbital | Frontal_Med_Orb_L | 83.87 | Frontal_Med_Orb_R | 89.14 |
| Gyrus rectus | Rectus_L | 84.04 | Rectus_R | 88.72 |
| Medial orbital gyrus | OFCmed_L | 73.27 | OFCmed_R | 66.02 |
| Anterior orbital gyrus | OFCant_L | 73.36 | OFCant_R | 65.59 |
| Posterior orbital gyrus | OFCpost_L | 67.90 | OFCpost_R | 57.40 |
| Lateral orbital gyrus | OFClat_L | 85.28 | - | - |
| Insula | Insula_L | 77.56 | Insula_R | 81.30 |
| Hippocampus | Hippocampus_L | 72.32 | Hippocampus_R | 76.00 |
| Parahippocampal gyrus | ParaHippocampal_L | 55.62 | ParaHippocampal_R | 50.35 |
| Amygdala | Amygdala_L | 87.27 | Amygdala_R | 77.02 |
| Anterior cingulate cortex, subgenual | ACC_sub_L | 94.64 | ACC_sub_R | 97.73 |
| Anterior cingulate cortex, pregenual | ACC_pre_L | 96.17 | ACC_pre_R | 92.90 |
| Anterior cingulate cortex, supracallosal | ACC_sup_L | 88.43 | ACC_sup_R | 90.81 |

**Table S3.** Complete summary of linear mixed effects model (LMM) with linear terms (target emotion: sadness).

| **Model equation:** | **\$metric-score\$ ~ time + (1\|Participant)** | | |
|---|---|---|---|
| Method: | REML | No. Groups: | 14 |
| No. Observations: | 812 | Group size: | 58 |

**Dependent Variable: correlation**

| Model Converged: | Yes | Scale: | 0.0161 | Log-Likelihood: | 475.4604 |
|---|---|---|---|---|---|
| **Fixed Effects** | | | | | |
| | **Coef.** | **Std.Err.** | **z** | **P>\|z\|** | **95% CI** |
| Intercept | 0.139 | 0.080 | 1.733 | 0.083 | -0.018 ~ 0.296 |
| time | 0.003 | 0.000 | 9.438 | < 0.001 | 0.002 ~ 0.003 |
| **Random Effects (Group Var)** | | | | | |
| Coef.: | 0.089 | Std.Err.: | 0.276 | | |

**Dependent Variable: angular_distance**

| Model Converged: | Yes | Scale: | 488.3123 | Log-Likelihood: | -3697.7605 |
|---|---|---|---|---|---|
| **Fixed Effects** | | | | | |
| | **Coef.** | **Std.Err.** | **z** | **P>\|z\|** | **95% CI** |
| Intercept | 75.907 | 8.913 | 8.517 | < 0.001 | 58.438 ~ 93.375 |
| time | -0.447 | 0.046 | -9.644 | < 0.001 | -0.538 ~ -0.356 |
| **Random Effects (Group Var)** | | | | | |
| Coef.: | 1073.871 | Std.Err.: | 19.358 | | |

**Dependent Variable: intensity**

| Model Converged: | Yes | Scale: | 0.7921 | Log-Likelihood: | -1095.8233 |
|---|---|---|---|---|---|
| **Fixed Effects** | | | | | |
| | **Coef.** | **Std.Err.** | **z** | **P>\|z\|** | **95% CI** |
| Intercept | 0.658 | 0.353 | 1.862 | 0.063 | -0.034 ~ 1.350 |
| time | 0.020 | 0.002 | 10.709 | < 0.001 | 0.016 ~ 0.024 |
| **Random Effects (Group Var)** | | | | | |
| Coef.: | 1.683 | Std.Err.: | 0.754 | | |

**Table S4.** Likelihood ratio test (LRT) results for linear mixed-effects model comparison between quadratic or exponential model with linear model (target emotion: sadness).

**Model fit**

| | | | | | |
|---|---|---|---|---|---|
| Method: | ML | No. Runs: | 1 | | |
| No. Observations: | 812 | No. Groups: | 14 | Group size: | 58 |

**Dependent Variable: correlation**

| Model | Fixed Effects | Random Effects | AIC | BIC | LL | df |
|---|---|---|---|---|---|---|
| linear | time | Group (Intercept) | -960.81 | -942.01 | 484.40 | 3 |
| exponential | time + $e^{-time}$ | Group (Intercept) | -1045.17 | -1021.67 | 527.58 | 4 |
| quadratic | time + $time^2$ | Group (Intercept) | -1017.16 | -993.66 | 513.58 | 4 |

**Likelihood ratio test**

| | | | | | | |
|---|---|---|---|---|---|---|
| exponential vs. linear | df difference: | 1 | $\chi^2$: | 86.36 | p-value: | 1.50e-20 |
| quadratic vs. linear | df difference: | 1 | $\chi^2$: | 58.36 | p-value: | 2.19e-14 |

**Dependent Variable: angular_distance**

| Model | Fixed Effects | Random Effects | AIC | BIC | LL | df |
|---|---|---|---|---|---|---|
| linear | time | Group (Intercept) | 7405.36 | 7424.16 | -3698.68 | 3 |
| exponential | time + $e^{-time}$ | Group (Intercept) | 7397.10 | 7420.60 | -3693.55 | 4 |
| quadratic | time + $time^2$ | Group (Intercept) | 7396.31 | 7419.80 | -3693.15 | 4 |

**Likelihood ratio test**

| | | | | | | |
|---|---|---|---|---|---|---|
| exponential vs. linear | df difference: | 1 | $\chi^2$: | 10.26 | p-value: | 0.00136 |
| quadratic vs. linear | df difference: | 1 | $\chi^2$: | 11.06 | p-value: | 8.83e-4 |

**Dependent Variable: intensity**

| Model | Fixed Effects | Random Effects | AIC | BIC | LL | df |
|---|---|---|---|---|---|---|
| linear | time | Group (Intercept) | 2188.61 | 2207.40 | -1090.30 | 3 |
| exponential | time + $e^{-time}$ | Group (Intercept) | 2162.54 | 2186.04 | -1076.27 | 4 |
| quadratic | time + $time^2$ | Group (Intercept) | 2182.45 | 2205.94 | -1086.22 | 4 |

**Likelihood ratio test**

| | | | | | | |
|---|---|---|---|---|---|---|
| exponential vs. linear | df difference: | 1 | $\chi^2$: | 28.06 | p-value: | 1.17e-07 |
| quadratic vs. linear | df difference: | 1 | $\chi^2$: | 8.16 | p-value: | 0.00428 |

**Table S5.** Complete summary of linear mixed effects model (LMM) with quadratic terms (target emotion: sadness).

| **Model equation:** | **\$metric-score\$ ~ time + $time^2$ + (1\|Participant)** | | |
|---|---|---|---|
| Method: | REML | No. Groups: | 14 |
| No. Observations: | 812 | Group size: | 58 |

**Dependent Variable: correlation**

| Model Converged: | Yes | Scale: | 0.0145 | Log-Likelihood: | 508.5160 |
|---|---|---|---|---|---|
| **Fixed Effects** | | | | | |
| | **Coef.** | **Std.Err.** | **z** | **P>\|z\|** | **95% CI** |
| Intercept | 0.024 | 0.081 | 0.301 | 0.764 | -0.134 ~ 0.183 |
| time | 0.013 | 0.001 | 11.581 | < 0.001 | 0.011 ~ 0.015 |
| $time^2$ | -0.000 | 0.000 | -9.538 | < 0.001 | -0.000 ~ -0.000 |
| **Random Effects (Group Var)** | | | | | |
| Coef.: | 0.089 | Std.Err.: | 0.292 | | |

**Dependent Variable: angular_distance**

| Model Converged: | Yes | Scale: | 482.6797 | Log-Likelihood: | -3697.5016 |
|---|---|---|---|---|---|
| **Fixed Effects** | | | | | |
| | **Coef.** | **Std.Err.** | **z** | **P>\|z\|** | **95% CI** |
| Intercept | 82.938 | 9.176 | 9.038 | < 0.001 | 64.953 ~ 100.924 |
| time | -1.069 | 0.199 | -5.365 | < 0.001 | -1.460 ~ -0.678 |
| $time^2$ | 0.010 | 0.003 | 3.210 | 0.001 | 0.004 ~ 0.016 |
| **Random Effects (Group Var)** | | | | | |
| Coef.: | 1073.913 | Std.Err.: | 19.469 | | |

**Dependent Variable: intensity**

| Model Converged: | Yes | Scale: | 0.7657 | Log-Likelihood: | -1089.8969 |
|---|---|---|---|---|---|
| **Fixed Effects** | | | | | |
| | **Coef.** | **Std.Err.** | **z** | **P>\|z\|** | **95% CI** |
| Intercept | 0.192 | 0.364 | 0.528 | 0.598 | -0.521 ~ 0.904 |
| time | 0.061 | 0.008 | 7.711 | < 0.001 | 0.046 ~ 0.077 |
| $time^2$ | -0.001 | 0.000 | -5.338 | < 0.001 | -0.001 ~ -0.000 |
| **Random Effects (Group Var)** | | | | | |
| Coef.: | 1.684 | Std.Err.: | 0.766 | | |

**Table S6.** Complete summary of linear mixed effects model (LMM) with exponential terms (target emotion: sadness).

| **Model equation:** | **\$metric-score\$ ~ time + $e^{time}$ + (1\|Participant)** | | |
|---|---|---|---|
| Method: | REML | No. Groups: | 14 |
| No. Observations: | 812 | Group size: | 58 |

**Dependent Variable: correlation**

| Model Converged: | Yes | Scale: | 0.0150 | Log-Likelihood: | 505.1043 |
|---|---|---|---|---|---|
| | | **Fixed Effects** | | | |
| | **Coef.** | **Std.Err.** | **z** | **P>\|z\|** | **95% CI** |
| Intercept | 0.168 | 0.080 | 2.092 | 0.036 | 0.011 ~ 0.325 |
| time | 0.002 | 0.000 | 6.689 | < 0.001 | 0.001 ~ 0.002 |
| $e^{time}$ | -5.126 | 0.660 | -7.771 | < 0.001 | -6.419 ~ -3.833 |
| | | **Random Effects (Group Var)** | | | |
| Coef.: | 0.089 | Std.Err.: | 0.286 | | |

**Dependent Variable: angular_distance**

| Model Converged: | Yes | Scale: | 482.1982 | Log-Likelihood: | -3686.5472 |
|---|---|---|---|---|---|
| | | **Fixed Effects** | | | |
| | **Coef.** | **Std.Err.** | **z** | **P>\|z\|** | **95% CI** |
| Intercept | 73.692 | 8.936 | 8.247 | < 0.001 | 56.178 ~ 91.205 |
| time | -0.393 | 0.049 | -8.073 | < 0.001 | -0.489 ~ -0.298 |
| $e^{time}$ | 394.150 | 118.271 | 3.333 | 0.001 | 162.343 ~ 625.956 |
| | | **Random Effects (Group Var)** | | | |
| Coef.: | 1073.916 | Std.Err.: | 19.479 | | |

**Dependent Variable: intensity**

| Model Converged: | Yes | Scale: | 0.7851 | Log-Likelihood: | -1089.2671 |
|---|---|---|---|---|---|
| | | **Fixed Effects** | | | |
| | **Coef.** | **Std.Err.** | **z** | **P>\|z\|** | **95% CI** |
| Intercept | 0.734 | 0.354 | 2.074 | 0.038 | 0.040 ~ 1.428 |
| time | 0.018 | 0.002 | 9.221 | 0.000 | 0.014 ~ 0.022 |
| $e^{time}$ | -13.650 | 4.722 | -2.860 | 0.004 | -23.003 ~ -4.296 |
| | | **Random Effects (Group Var)** | | | |
| Coef.: | 1.684 | Std.Err.: | 0.757 | | |

**Table S7.** Individual feedback performance for sadness (N = 14), assessed using linear regression across time (excluding the first two time points). The “success” column indicates whether the corresponding metric showed a significant improvement (p < 0.05, FDR-corrected). Specifically, success was defined as a significantly positive slope for Pearson’s correlation and intensity level, and a significantly negative slope for angular distance.

| sub_id | run | Pearson’s correlation | | | angular distance | | | intensity level | | |
|---|---|---|---|---|---|---|---|---|---|---|
| | | slope | p_value | success | slope | p_value | success | slope | p_value | success |
| S01 | Run 1 | 0.001 | 0.39 | N | -0.212 | < 0.001 | Y | 0.06 | < 0.001 | Y |
| S02 | Run 1 | 0.001 | 0.232 | N | 0.015 | 0.602 | N | -0.02 | 0.013 | N |
| S03 | Run 1 | 0.002 | 0.002 | Y | -0.624 | 0.089 | N | 0 | 1 | N |
| S08 | Run 1 | 0.005 | < 0.001 | Y | -2.008 | < 0.001 | Y | 0 | 1 | N |
| S09 | Run 1 | 0.01 | < 0.001 | Y | -0.783 | < 0.001 | Y | 0.095 | < 0.001 | Y |
| S10 | Run 1 | -0.003 | < 0.001 | N | 0.147 | 0.251 | N | 0 | 1 | N |
| S11 | Run 1 | 0 | 0.667 | N | 0.01 | 0.602 | N | 0 | 1 | N |
| S12 | Run 1 | 0.001 | 0.139 | N | -0.23 | 0.089 | N | -0.003 | 0.069 | N |
| S14 | Run 1 | 0.001 | 0.121 | N | -0.073 | < 0.001 | Y | 0.078 | < 0.001 | Y |
| S16 | Run 1 | 0.004 | 0.013 | Y | -0.15 | 0.342 | N | -0.014 | 0.013 | N |
| S17 | Run 1 | 0.01 | < 0.001 | Y | -1.547 | < 0.001 | Y | 0.04 | < 0.001 | Y |
| S19 | Run 1 | 0.003 | < 0.001 | Y | -0.843 | < 0.001 | Y | 0.047 | < 0.001 | Y |
| S21 | Run 1 | 0.003 | 0.007 | Y | -0.116 | 0.001 | Y | -0.008 | 0.451 | N |
| S25 | Run 1 | -0.002 | 0.015 | N | 0.159 | < 0.001 | N | 0.005 | 0.291 | N |

**Table S8.** Complete summary of linear mixed effects model (LMM) with linear terms (target emotion: joyful relaxation).

| **Model equation:** | $metric-score$ ~ time + run + time:run + (1\|Participant) | | |
|---|---|---|---|
| Method: | REML | No. Groups: | 24 |
| No. Observations: | 2784 | Group size: | 116 |

**Dependent Variable: correlation**

| Model Converged: | Yes | Scale: | 0.0279 | Log-Likelihood: | 949.5174 |
|---|---|---|---|---|---|
| **Fixed Effects** | | | | | |
| | **Coef.** | **Std.Err.** | **z** | **P>\|z\|** | **95% CI** |
| Intercept [Run 1] | 0.061 | 0.050 | 1.236 | 0.217 | -0.036 ~ 0.158 |
| Intercept [Run 2] | 0.111 | 0.050 | 2.245 | 0.025 | 0.014 ~ 0.208 |
| Intercept [Run 2 vs. Run 1] | 0.050 | 0.013 | 3.710 | < 0.001 | 0.024 ~ 0.076 |
| time [Run 1] | 0.002 | < 0.001 | 7.432 | < 0.001 | 0.001 ~ 0.003 |
| time [Run 2] | 0.002 | < 0.001 | 6.431 | < 0.001 | 0.001 ~ 0.002 |
| time [Run 2 vs. Run 1] | > -0.001 | < 0.001 | -0.708 | 0.479 | -0.001 ~ < 0.001 |
| **Random Effects (Group Var)** | | | | | |
| Coef.: | 0.057 | Std.Err.: | 0.101 | | |

**Dependent Variable: angular_distance**

| Model Converged: | Yes | Scale: | 768.8770 | Log-Likelihood: | -13260.4152 |
|---|---|---|---|---|---|
| **Fixed Effects** | | | | | |
| | **Coef.** | **Std.Err.** | **z** | **P>\|z\|** | **95% CI** |
| Intercept [Run 1] | 81.819 | 7.053 | 11.600 | < 0.001 | 67.995 ~ 95.643 |
| Intercept [Run 2] | 77.177 | 7.053 | 10.942 | < 0.001 | 63.353 ~ 91.001 |
| Intercept [Run 2 vs. Run 1] | -4.642 | 2.240 | -2.073 | 0.038 | -9.032 ~ -0.252 |
| time [Run 1] | -0.172 | 0.044 | -3.863 | < 0.001 | -0.259 ~ -0.084 |
| time [Run 2] | -0.232 | 0.044 | -5.221 | < 0.001 | -0.319 ~ -0.145 |
| time [Run 2 vs. Run 1] | -0.060 | 0.063 | -0.960 | 0.337 | -0.183 ~ 0.063 |
| **Random Effects (Group Var)** | | | | | |
| Coef.: | 1133.735 | Std.Err.: | 12.177 | | |

**Dependent Variable: intensity**

| Model Converged: | Yes | Scale: | 1.0905 | Log-Likelihood: | -4146.6694 |
|---|---|---|---|---|---|
| **Fixed Effects** | | | | | |
| | **Coef.** | **Std.Err.** | **z** | **P>\|z\|** | **95% CI** |
| Intercept [Run 1] | 0.525 | 0.293 | 1.794 | 0.073 | -0.049 ~ 1.099 |
| Intercept [Run 2] | 1.025 | 0.293 | 3.499 | < 0.001 | 0.451 ~ 1.599 |
| Intercept [Run 2 vs. Run 1] | 0.499 | 0.084 | 5.921 | < 0.001 | 0.334 ~ 0.665 |
| time [Run 1] | 0.017 | 0.002 | 9.995 | < 0.001 | 0.013 ~ 0.020 |
| time [Run 2] | 0.016 | 0.002 | 9.447 | < 0.001 | 0.013 ~ 0.019 |
| time [Run 2 vs. Run 1] | -0.001 | 0.002 | -0.388 | 0.698 | -0.006 ~ 0.004 |
| **Random Effects (Group Var)** | | | | | |
| Coef.: | 1.974 | Std.Err.: | 0.562 | | |

**Table S9.** Likelihood ratio test (LRT) results for linear mixed-effects model comparison between quadratic or exponential model with linear model (target emotion: joyful relaxation).

**Model fit**

| | | | | | |
|---|---|---|---|---|---|
| Method: | ML | No. Runs: | 2 | | |
| No. Observations: | 2784 | No. Groups: | 24 | Group size: | 116 |

**Dependent Variable: correlation**

| Model | Fixed Effects | Random Effects | AIC | BIC | LL | df |
|---|---|---|---|---|---|---|
| linear | time + run + time:run | Group (Intercept) | -1928.79 | -1893.20 | 970.39 | 5 |
| exponential | time + run + time:run + $e^{-time}$ + $e^{-time}$:run | Group (Intercept) | -1931.82 | -1884.37 | 973.91 | 7 |
| quadratic | time + run + time:run + $time^2$ + $time^2$:run | Group (Intercept) | -1939.03 | -1891.58 | 977.52 | 7 |

**Likelihood ratio test**

| | | | | | | |
|---|---|---|---|---|---|---|
| exponential vs. linear | df difference: | 2 | $\chi^2$: | 7.03 | p-value: | 0.0297 |
| quadratic vs. linear | df difference: | 2 | $\chi^2$: | 14.25 | p-value: | 8.05e-04 |

**Dependent Variable: angular_distance**

| Model | Fixed Effects | Random Effects | AIC | BIC | LL | df |
|---|---|---|---|---|---|---|
| linear | time + run + time:run | Group (Intercept) | 26531.66 | 26567.25 | -13259.83 | 5 |
| exponential | time + run + time:run + $e^{-time}$ + $e^{-time}$:run | Group (Intercept) | 26522.71 | 26570.16 | -13253.36 | 7 |
| quadratic | time + run + time:run + $time^2$ + $time^2$:run | Group (Intercept) | 26528.56 | 26576.01 | -13256.28 | 7 |

**Likelihood ratio test**

| | | | | | | |
|---|---|---|---|---|---|---|
| exponential vs. linear | df difference: | 2 | $\chi^2$: | 12.95 | p-value: | 0.00154 |
| quadratic vs. linear | df difference: | 2 | $\chi^2$: | 7.10 | p-value: | 0.0287 |

**Dependent Variable: intensity**

| Model | Fixed Effects | Random Effects | AIC | BIC | LL | df |
|---|---|---|---|---|---|---|
| linear | time + run + time:run | Group (Intercept) | 8278.14 | 8313.73 | -4133.07 | 5 |
| exponential | time + run + time:run + $e^{-time}$ + $e^{-time}$:run | Group (Intercept) | 8260.69 | 8308.15 | -4122.35 | 7 |
| quadratic | time + run + time:run + $time^2$ + $time^2$:run | Group (Intercept) | 8212.03 | 8259.48 | -4098.01 | 7 |

**Likelihood ratio test**

| | | | | | | |
|---|---|---|---|---|---|---|
| exponential vs. linear | df difference: | 2 | $\chi^2$: | 21.45 | p-value: | 2.20e-05 |
| quadratic vs. linear | df difference: | 2 | $\chi^2$: | 70.12 | p-value: | 5.95e-16 |

**Table S10.** Complete summary of linear mixed effects model (LMM) with quadratic terms (target emotion: joyful relaxation).

| **Model equation:** | **\$metric-score\$ ~ time + run + time:run + time$^2$ + time$^2$:run + (1\|Participant)** | | |
|---|---|---|---|
| Method: | REML | No. Groups: | 24 |
| No. Observations: | 2784 | Group size: | 116 |

**Dependent Variable: correlation**

| Model Converged: | Yes | Scale: | 0.0277 | Log-Likelihood: | 936.6004 |
|---|---|---|---|---|---|
| **Fixed Effects** | | | | | |
| | **Coef.** | **Std.Err.** | **z** | **P>\|z\|** | **95% CI** |
| Intercept [Run 1] | 0.060 | 0.051 | 1.175 | 0.240 | -0.040 ~ 0.160 |
| Intercept [Run 2] | 0.063 | 0.051 | 1.239 | 0.215 | -0.037 ~ 0.164 |
| Intercept [Run 2 vs. Run 1] | 0.003 | 0.022 | 0.145 | 0.885 | -0.041 ~ 0.047 |
| time [Run 1] | 0.002 | 0.001 | 1.808 | 0.071 | > -0.001 ~ 0.004 |
| time [Run 2] | 0.006 | 0.001 | 5.163 | < 0.001 | 0.004 ~ 0.008 |
| time [Run 2 vs. Run 1] | 0.004 | 0.002 | 2.372 | 0.018 | 0.001 ~ 0.007 |
| time$^2$ [Run 1] | > -0.001 | < 0.001 | -0.088 | 0.930 | > -0.001 ~ < 0.001 |
| time$^2$ [Run 2] | > -0.001 | < 0.001 | -3.775 | < 0.001 | > -0.001 ~ > -0.001 |
| time$^2$ [Run 2 vs. Run 1] | > -0.001 | < 0.001 | -2.607 | 0.009 | > -0.001 ~ > -0.001 |
| **Random Effects (Group Var)** | | | | | |
| Coef.: | 0.057 | Std.Err.: | 0.101 | | |

**Dependent Variable: angular_distance**

| Model Converged: | Yes | Scale: | 767.4573 | Log-Likelihood: | -13266.6724 |
|---|---|---|---|---|---|
| **Fixed Effects** | | | | | |
| | **Coef.** | **Std.Err.** | **z** | **P>\|z\|** | **95% CI** |
| Intercept [Run 1] | 81.099 | 7.362 | 11.016 | < 0.001 | 66.671 ~ 95.528 |
| Intercept [Run 2] | 82.753 | 7.362 | 11.241 | < 0.001 | 68.324 ~ 97.182 |
| Intercept [Run 2 vs. Run 1] | 1.654 | 3.730 | 0.443 | 0.657 | -5.656 ~ 8.964 |
| time [Run 1] | -0.108 | 0.192 | -0.562 | 0.574 | -0.484 ~ 0.268 |
| time [Run 2] | -0.725 | 0.192 | -3.779 | < 0.001 | -1.101 ~ -0.349 |
| time [Run 2 vs. Run 1] | -0.617 | 0.271 | -2.275 | 0.023 | -1.149 ~ -0.085 |
| time$^2$ [Run 1] | -0.001 | 0.003 | -0.341 | 0.733 | -0.007 ~ 0.005 |
| time$^2$ [Run 2] | 0.008 | 0.003 | 2.643 | 0.008 | 0.002 ~ 0.014 |
| time$^2$ [Run 2 vs. Run 1] | 0.009 | 0.004 | 2.110 | 0.035 | 0.001 ~ 0.017 |
| **Random Effects (Group Var)** | | | | | |
| Coef.: | 1133.746 | Std.Err.: | 12.188 | | |

**Dependent Variable: intensity**

| Model Converged: | Yes | Scale: | 1.0639 | Log-Likelihood: | -4128.0362 |
|---|---|---|---|---|---|
| **Fixed Effects** | | | | | |
| | **Coef.** | **Std.Err.** | **z** | **P>\|z\|** | **95% CI** |
| Intercept [Run 1] | 0.583 | 0.303 | 1.922 | 0.055 | -0.011 ~ 1.177 |
| Intercept [Run 2] | 0.366 | 0.303 | 1.207 | 0.227 | -0.228 ~ 0.960 |
| Intercept [Run 2 vs. Run 1] | -0.217 | 0.139 | -1.562 | 0.118 | -0.489 ~ 0.055 |
| time [Run 1] | 0.012 | 0.007 | 1.629 | 0.103 | -0.002 ~ 0.026 |
| time [Run 2] | 0.074 | 0.007 | 10.371 | < 0.001 | 0.060 ~ 0.088 |
| time [Run 2 vs. Run 1] | 0.062 | 0.010 | 6.182 | < 0.001 | 0.043 ~ 0.082 |
| time$^2$ [Run 1] | < 0.001 | < 0.001 | 0.730 | 0.465 | > -0.001 ~ < 0.001 |
| time$^2$ [Run 2] | -0.001 | < 0.001 | -8.388 | < 0.001 | -0.001 ~ -0.001 |
| time$^2$ [Run 2 vs. Run 1] | -0.001 | < 0.001 | -6.447 | < 0.001 | -0.001 ~ -0.001 |
| **Random Effects (Group Var)** | | | | | |
| Coef.: | 1.974 | Std.Err.: | 0.569 | | |

**Table S11.** Complete summary of linear mixed effects model (LMM) with exponential terms (target emotion: joyful relaxation).

| **Model equation:** | **\$metric-score\$ ~ time + run + time:run + $e^{time}$ + $e^{time}$:run + (1\|Participant)** | | |
|---|---|---|---|
| Method: | REML | No. Groups: | 24 |
| No. Observations: | 2784 | Group size: | 116 |

**Dependent Variable: correlation**

| Model Converged: | Yes | Scale: | 0.0278 | Log-Likelihood: | 954.1143 |
|---|---|---|---|---|---|
| **Fixed Effects** | | | | | |
| | **Coef.** | **Std.Err.** | **z** | **P>\|z\|** | **95% CI** |
| Intercept [Run 1] | 0.061 | 0.050 | 1.233 | 0.217 | -0.036 ~ 0.159 |
| Intercept [Run 2] | 0.121 | 0.050 | 2.444 | 0.015 | 0.024 ~ 0.219 |
| Intercept [Run 2 vs. Run 1] | 0.060 | 0.015 | 4.141 | < 0.001 | 0.032 ~ 0.089 |
| time [Run 1] | 0.002 | < 0.001 | 7.021 | < 0.001 | 0.001 ~ 0.003 |
| time [Run 2] | 0.001 | < 0.001 | 5.210 | < 0.001 | 0.001 ~ 0.002 |
| time [Run 2 vs. Run 1] | -0.001 | < 0.001 | -1.281 | 0.200 | -0.001 ~ < 0.001 |
| $e^{time}$ [Run 1] | -0.011 | 0.686 | -0.017 | 0.987 | -1.356 ~ 1.333 |
| $e^{time}$ [Run 2] | -1.819 | 0.686 | -2.651 | 0.008 | -3.163 ~ -0.474 |
| $e^{time}$ [Run 2 vs. Run 1] | -1.807 | 0.970 | -1.863 | 0.062 | -3.709 ~ 0.094 |
| **Random Effects (Group Var)** | | | | | |
| Coef.: | 0.057 | Std.Err.: | 0.101 | | |

**Dependent Variable: angular_distance**

| Model Converged: | Yes | Scale: | 765.8333 | Log-Likelihood: | -13242.6399 |
|---|---|---|---|---|---|
| **Fixed Effects** | | | | | |
| | **Coef.** | **Std.Err.** | **z** | **P>\|z\|** | **95% CI** |
| Intercept [Run 1] | 81.091 | 7.081 | 11.451 | < 0.001 | 67.211 ~ 94.970 |
| Intercept [Run 2] | 74.993 | 7.081 | 10.590 | < 0.001 | 61.113 ~ 88.872 |
| Intercept [Run 2 vs. Run 1] | -6.098 | 2.411 | -2.529 | 0.011 | -10.824 ~ -1.372 |
| time [Run 1] | -0.154 | 0.047 | -3.282 | 0.001 | -0.246 ~ -0.062 |
| time [Run 2] | -0.179 | 0.047 | -3.821 | < 0.001 | -0.271 ~ -0.087 |
| time [Run 2 vs. Run 1] | -0.025 | 0.066 | -0.381 | 0.703 | -0.155 ~ 0.105 |
| $e^{time}$ [Run 1] | 129.619 | 113.839 | 1.139 | 0.255 | -93.501 ~ 352.740 |
| $e^{time}$ [Run 2] | 388.738 | 113.839 | 3.415 | 0.001 | 165.618 ~ 611.859 |
| $e^{time}$ [Run 2 vs. Run 1] | 259.119 | 160.992 | 1.610 | 0.108 | -56.420 ~ 574.659 |
| **Random Effects (Group Var)** | | | | | |
| Coef.: | 1133.759 | Std.Err.: | 12.201 | | |

**Dependent Variable: intensity**

| Model Converged: | Yes | Scale: | 1.0828 | Log-Likelihood: | -4131.2125 |
|---|---|---|---|---|---|
| **Fixed Effects** | | | | | |
| | **Coef.** | **Std.Err.** | **z** | **P>\|z\|** | **95% CI** |
| Intercept [Run 1] | 0.541 | 0.294 | 1.839 | 0.066 | -0.035 ~ 1.116 |
| Intercept [Run 2] | 1.135 | 0.294 | 3.863 | < 0.001 | 0.559 ~ 1.711 |
| Intercept [Run 2 vs. Run 1] | 0.595 | 0.091 | 6.559 | < 0.001 | 0.417 ~ 0.772 |
| time [Run 1] | 0.016 | 0.002 | 9.267 | < 0.001 | 0.013 ~ 0.020 |
| time [Run 2] | 0.013 | 0.002 | 7.447 | < 0.001 | 0.010 ~ 0.017 |
| time [Run 2 vs. Run 1] | -0.003 | 0.002 | -1.287 | 0.198 | -0.008 ~ 0.002 |
| $e^{time}$ [Run 1] | -2.704 | 4.281 | -0.632 | 0.528 | -11.094 ~ 5.686 |
| $e^{time}$ [Run 2] | -19.658 | 4.281 | -4.593 | < 0.001 | -28.048 ~ -11.269 |
| $e^{time}$ [Run 2 vs. Run 1] | -16.955 | 6.054 | -2.801 | 0.005 | -28.819 ~ -5.090 |
| **Random Effects (Group Var)** | | | | | |
| Coef.: | 1.974 | Std.Err.: | 0.564 | | |

**Table S12.** Individual feedback performance for joyful relaxation (N = 24), assessed using linear regression across time (excluding the first two time points). The “success” column indicates whether the corresponding metric showed a significant improvement ($p < 0.05$, FDR-corrected). Specifically, success was defined as a significantly positive slope for Pearson’s correlation and intensity level, and a significantly negative slope for angular distance.

| sub_id | run | Pearson’s correlation | | | angular distance | | | intensity level | | |
|---|---|---|---|---|---|---|---|---|---|---|
| | | slope | p_value | success | slope | p_value | success | slope | p_value | success |
| S02 | Run 1 | 0.012 | < 0.001 | Y | -0.532 | < 0.001 | Y | 0.067 | < 0.001 | Y |
| | Run 2 | 0 | 0.821 | N | 0.07 | < 0.001 | N | 0.02 | 0.005 | Y |
| S03 | Run 1 | 0 | 0.779 | N | 1.502 | < 0.001 | N | -0.031 | < 0.001 | N |
| | Run 2 | 0.008 | < 0.001 | Y | 0.636 | 0.001 | N | 0 | 1 | N |
| S04 | Run 1 | -0.002 | 0.035 | N | 1.035 | 0.006 | N | 0.002 | 1 | N |
| | Run 2 | 0.015 | < 0.001 | Y | -0.48 | 0.023 | Y | 0.058 | < 0.001 | Y |
| S05 | Run 1 | -0.007 | < 0.001 | N | 0.509 | < 0.001 | N | -0.006 | 0.008 | N |
| | Run 2 | 0.004 | < 0.001 | Y | 0.636 | 0.027 | N | -0.019 | < 0.001 | N |
| S06 | Run 1 | 0.003 | < 0.001 | Y | -1.749 | < 0.001 | Y | 0 | 1 | N |
| | Run 2 | -0.005 | < 0.001 | N | -2.445 | < 0.001 | Y | 0.043 | < 0.001 | Y |
| S07 | Run 1 | -0.001 | 0.382 | N | -0.338 | < 0.001 | Y | 0.029 | < 0.001 | Y |
| | Run 2 | -0.002 | 0.004 | N | -1.4 | < 0.001 | Y | 0.046 | < 0.001 | Y |
| S08 | Run 1 | 0.016 | < 0.001 | Y | -1.614 | < 0.001 | Y | 0.091 | < 0.001 | Y |
| | Run 2 | 0.008 | < 0.001 | Y | -0.171 | < 0.001 | Y | 0.047 | < 0.001 | Y |
| S09 | Run 1 | -0.006 | < 0.001 | N | 2.002 | < 0.001 | N | -0.008 | 0.04 | N |
| | Run 2 | -0.001 | 0.149 | N | -0.138 | < 0.001 | Y | 0 | 1 | N |
| S10 | Run 1 | -0.003 | < 0.001 | N | -0.167 | < 0.001 | Y | 0 | 1 | N |
| | Run 2 | 0.004 | < 0.001 | Y | 0.155 | < 0.001 | N | 0 | 1 | N |
| S11 | Run 1 | -0.001 | 0.357 | N | 0.087 | < 0.001 | N | -0.005 | 0.008 | N |
| | Run 2 | 0 | 0.969 | N | 0.027 | 0.123 | N | 0.004 | 0.716 | N |
| S12 | Run 1 | 0.008 | < 0.001 | Y | -1.077 | < 0.001 | Y | 0.026 | < 0.001 | Y |
| | Run 2 | 0.001 | 0.121 | N | -0.016 | 0.143 | N | 0.041 | < 0.001 | Y |
| S14 | Run 1 | 0.005 | < 0.001 | Y | 0.048 | 0.022 | N | 0.057 | < 0.001 | Y |
| | Run 2 | 0.003 | < 0.001 | Y | -0.282 | 0.023 | Y | 0.018 | 0.004 | Y |
| S15 | Run 1 | 0.004 | < 0.001 | Y | -1.136 | < 0.001 | Y | 0.01 | < 0.001 | Y |
| | Run 2 | 0 | 0.99 | N | 0.444 | 0.228 | N | -0.002 | 0.853 | N |
| S16 | Run 1 | 0.003 | < 0.001 | Y | 0.781 | < 0.001 | N | -0.011 | 0.002 | N |
| | Run 2 | 0 | 0.432 | N | 1.293 | < 0.001 | N | -0.012 | 0.013 | N |
| S17 | Run 1 | 0.002 | < 0.001 | Y | 0.442 | 0.012 | N | -0.014 | 0.291 | N |
| | Run 2 | 0.008 | < 0.001 | Y | 0.034 | 0.074 | N | 0.002 | 1 | N |
| S18 | Run 1 | 0.004 | < 0.001 | Y | 0.056 | 0.003 | N | 0 | 1 | N |
| | Run 2 | 0.002 | < 0.001 | Y | -0.17 | 0.606 | N | 0.017 | < 0.001 | Y |
| S19 | Run 1 | 0.003 | 0.077 | N | -0.741 | < 0.001 | Y | 0 | 1 | N |
| | Run 2 | 0.008 | < 0.001 | Y | -0.619 | 0.003 | Y | 0 | 1 | N |
| S20 | Run 1 | 0.002 | < 0.001 | Y | 0.04 | 0.275 | N | 0.035 | 0.024 | Y |
| | Run 2 | 0.004 | < 0.001 | Y | -1.68 | < 0.001 | Y | 0.067 | < 0.001 | Y |
| S21 | Run 1 | -0.004 | 0.015 | N | 0.153 | 0.137 | N | -0.003 | 0.224 | N |
| | Run 2 | -0.008 | < 0.001 | N | -0.163 | < 0.001 | Y | 0 | 1 | N |
| S22 | Run 1 | -0.008 | < 0.001 | N | 0.076 | < 0.001 | N | -0.007 | 0.037 | N |
| | Run 2 | -0.005 | < 0.001 | N | -0.018 | 0.219 | N | 0 | 1 | N |
| S23 | Run 1 | 0.003 | < 0.001 | Y | -0.186 | < 0.001 | Y | 0.061 | < 0.001 | Y |
| | Run 2 | -0.004 | 0.001 | N | -0.586 | 0.001 | Y | 0.034 | < 0.001 | Y |
| S25 | Run 1 | -0.001 | 0.117 | N | -1.875 | < 0.001 | Y | 0 | 1 | N |
| | Run 2 | -0.007 | < 0.001 | N | -0.762 | < 0.001 | Y | -0.029 | 0.034 | N |
| S26 | Run 1 | 0.011 | < 0.001 | Y | -0.04 | 0.131 | N | 0.057 | < 0.001 | Y |
| | Run 2 | 0.007 | < 0.001 | Y | -0.023 | 0.101 | N | 0.036 | < 0.001 | Y |
| S27 | Run 1 | 0.003 | 0.01 | Y | -1.392 | < 0.001 | Y | 0.05 | < 0.001 | Y |
| | Run 2 | 0 | 0.99 | N | 0.097 | 0.009 | N | 0.007 | 0.621 | N |

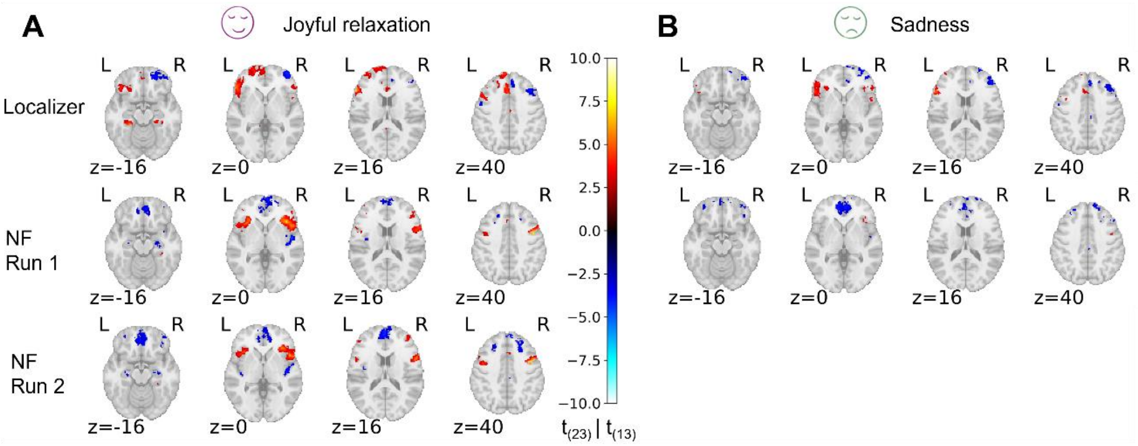


**Figure S1.** Group-level univariate analysis for NF runs and corresponding localizer runs targeting the same emotion. For consistency, univariate patterns for localizer runs were reconstructed using the same participant groups as in the respective NF runs. (A) Joyful relaxation (N = 24). (B) Sadness (N = 14). NF = neurofeedback.

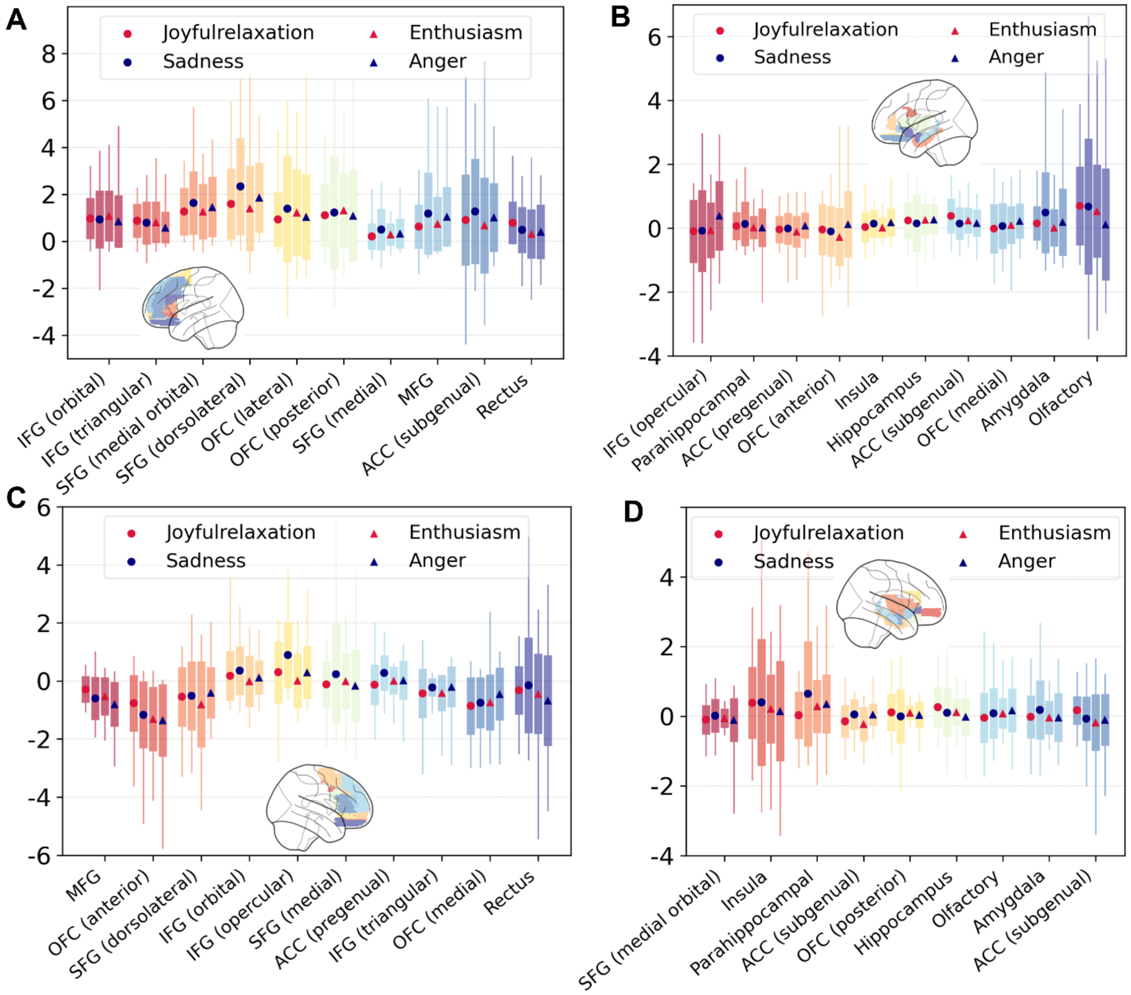


**Figure S2.** ROI analysis results across all 39 regions, based on the average activity of all voxels within each ROI. Regions are ordered by group-level activity across conditions for each hemisphere. (A) Top 10 regions in the left hemisphere. (B) Bottom 10 regions in the left hemisphere. (C) Top 10 regions in the right hemisphere. (D) Bottom 9 regions in the right hemisphere. ROI = region of interest; IFG = inferior frontal gyrus; SFG = superior frontal gyrus; OFC = orbitofrontal cortices; MFG = middle frontal gyrus; ACC = anterior cingulate cortex.

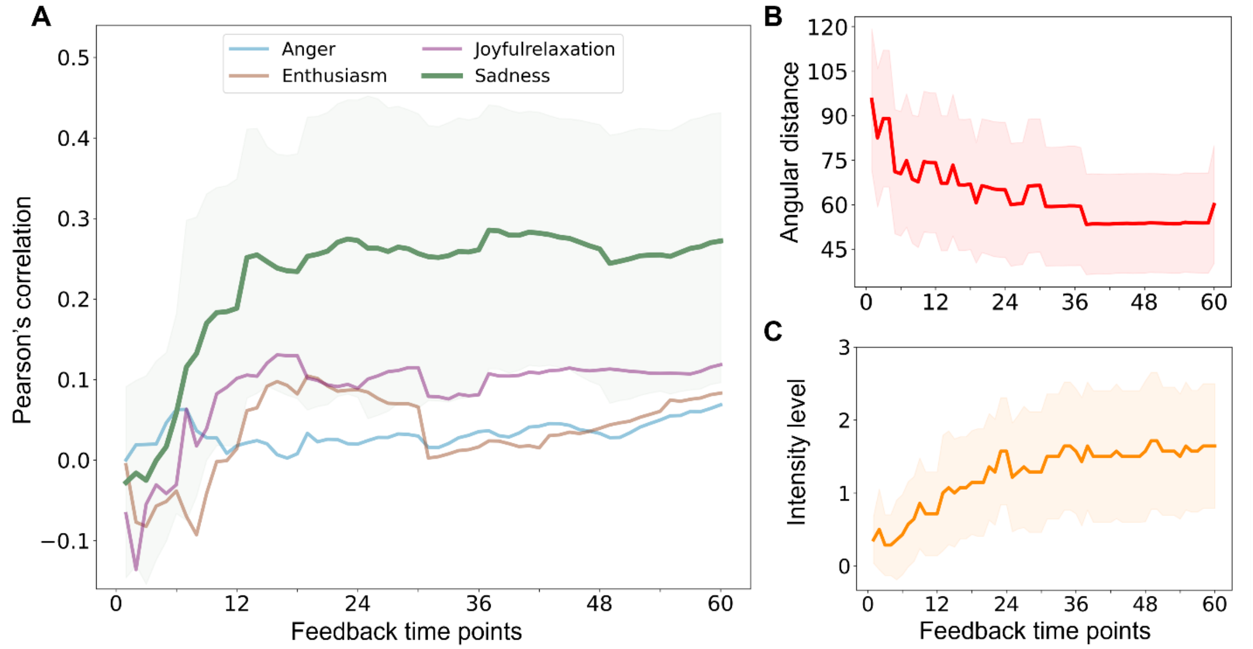


**Figure S3.** Learning curves of neurofeedback performance metrics (target condition: sadness). The metrics include Pearson's correlation (A), angular distance (B), and intensity level (C). In each part, the mean value (across participants) and its 95% confidence interval are plotted over all feedback time points for two runs. Part A additionally shows, in the background, Pearson's correlations between the real-time brain pattern and each non-target emotional base pattern.

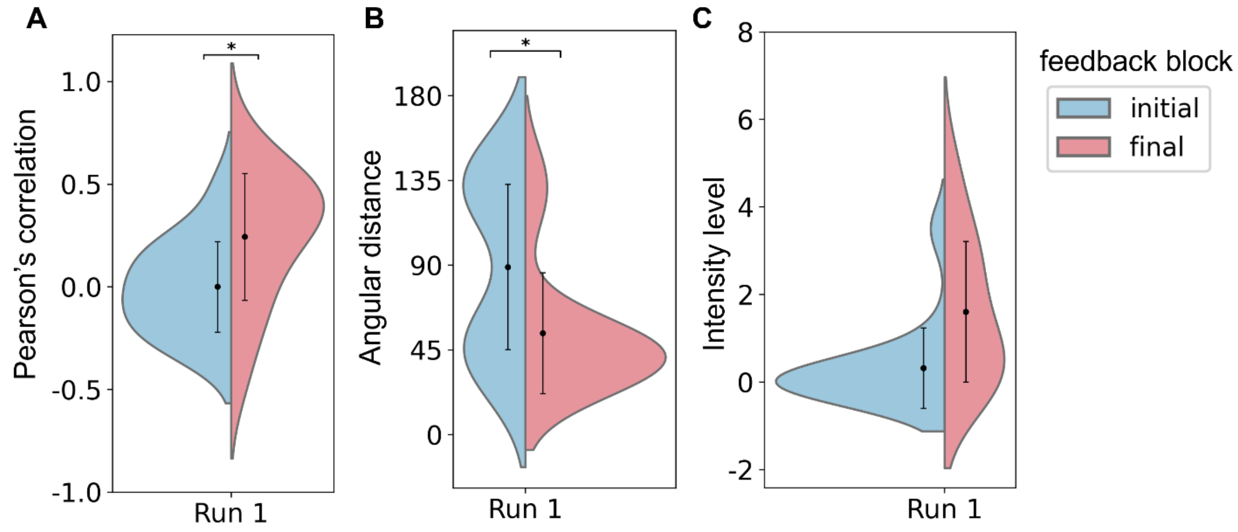


**Figure S4.** Comparison of neurofeedback performance metrics between initial and final feedback blocks (target condition: sadness). The median value of each metric within the corresponding block was extracted and compared. (A) Pearson's correlation coefficient between target pattern and the pattern extracted in the first and last feedback blocks. (B) Angular distance in the first and last feedback block, measured by calculating the angular distance between the red dot and the target stimulus on the CSM. (C) Same as (B) but using the thermometer reading of the target stimulus on the CSM as metric of intensity level. The violin plot shows the distribution of each metric across participants, and the error bar shows the corresponding mean and SD. Paired t-tests or Wilcoxon signed-rank tests were performed on the initial and final metrics. Results were considered significant (*: $p < 0.05$, **: $p < 0.01$) after Bonferroni correction ($n = 2$). CSM = circular semantic map; SD = standard deviation.

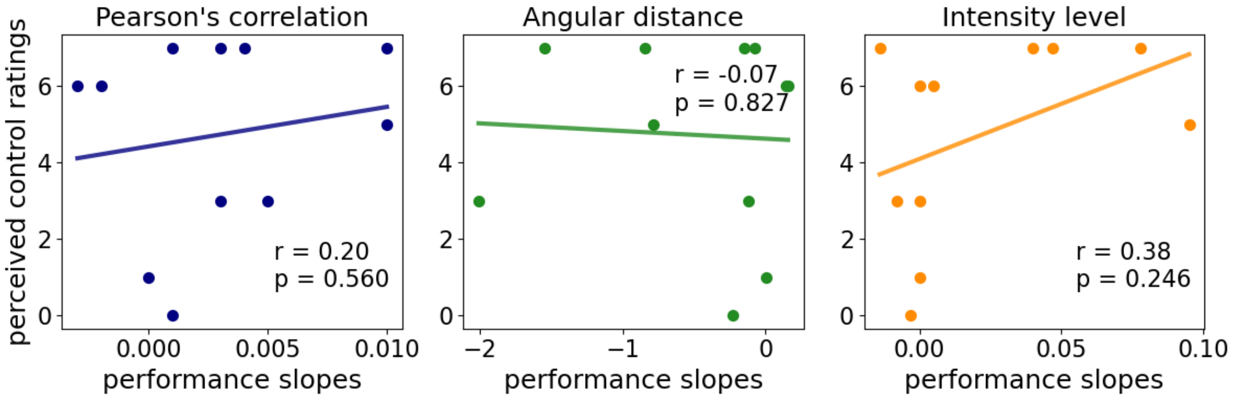


**Figure S5.** Correlation analyses between participants' perceived control ratings and neurofeedback performance slopes for sadness (N = 11). The perceived control ratings (y-axis) reflect participants' perceived control over brain activity, rated on a scale from 0 to 10. The performance slopes (x-axis) were obtained from individual linear regression models across the feedback time sequence. Statistical values were calculated using either Pearson's or Spearman's correlation (not corrected).

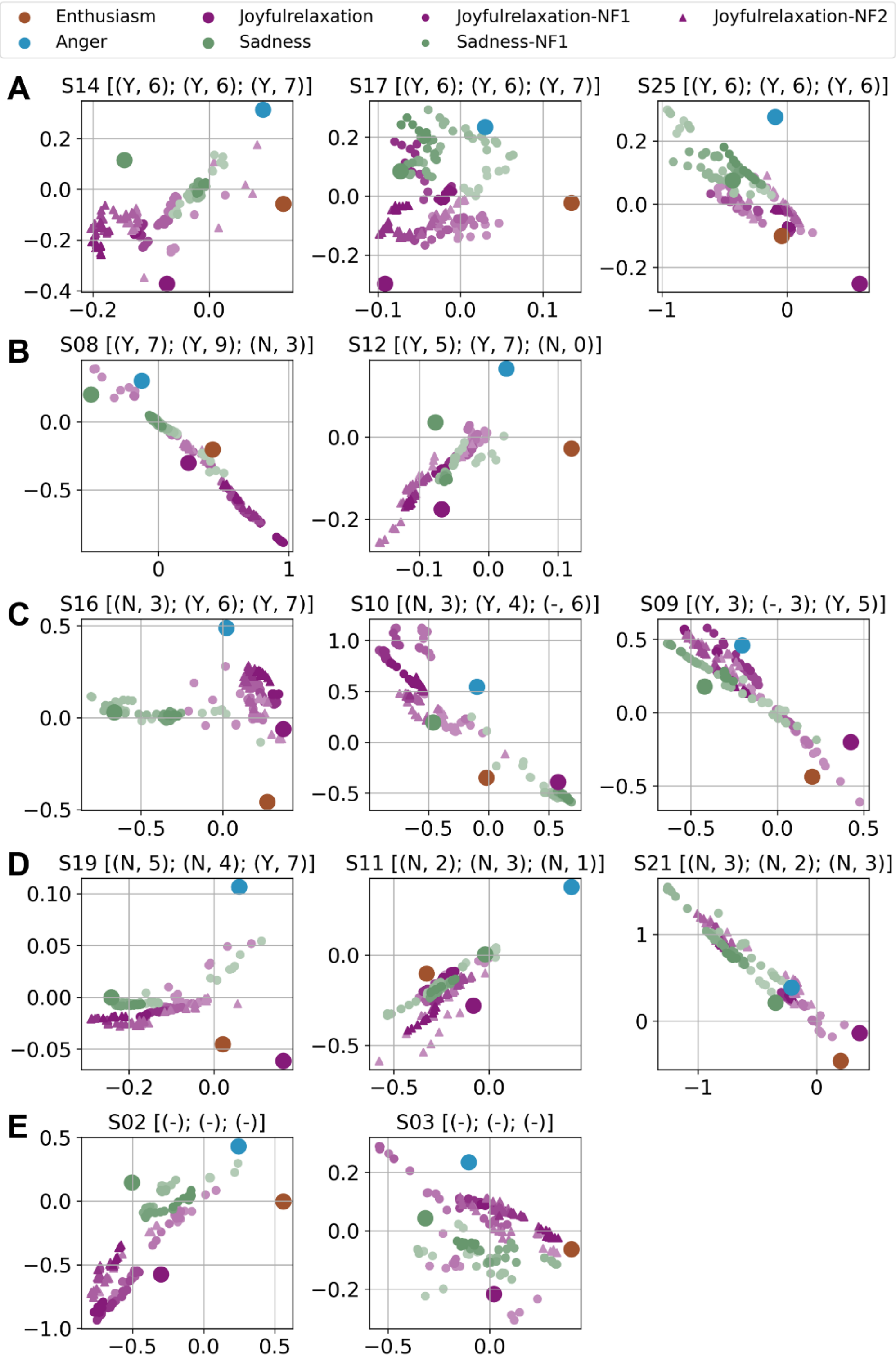


**Figure S6.** Individual modulation trajectories during neurofeedback runs visualized with MDS (for participants with both joyful relaxation and sadness runs, N = 13). The X and Y axes represent the two scaled dimensions from MDS. For each participant, the four localizer base patterns were reduced to two dimensions using MDS and plotted as anchor points (larger circles) with colors indicating the corresponding localizer emotion. Neurofeedback data were then projected into the same representational space for joyful relaxation (smaller circles for run 1 and smaller tringles for run 2) and for sadness (smaller circle) in the corresponding color, The shade of the makers indicates time progression, with darker shades showing later time points. Subfigure titles show the participant number and their perceived control ratings (Y/N = yes/no), followed by the reported rating score ('-' denotes no ratings collected or no corresponding run). Subpanels are arranged in descending order according to the number of positive ratings. MDS = multidimensional scaling.

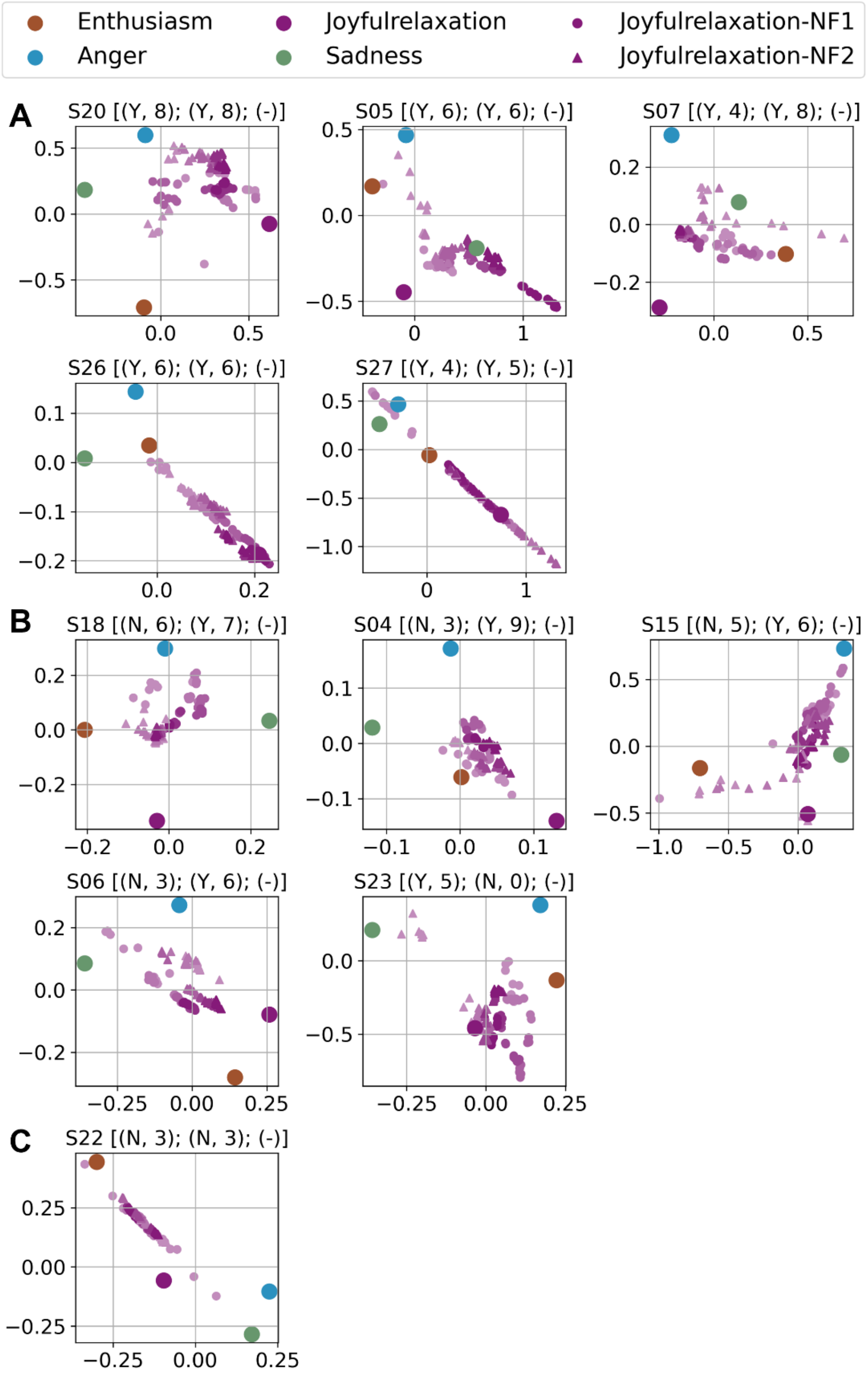


**Figure S7.** Individual modulation trajectories during neurofeedback runs visualized with MDS (for participants with only two joyful relaxation runs, N = 11). Processing procedure and plotting criteria are the same as in Figure S6. MDS = multidimensional scaling.

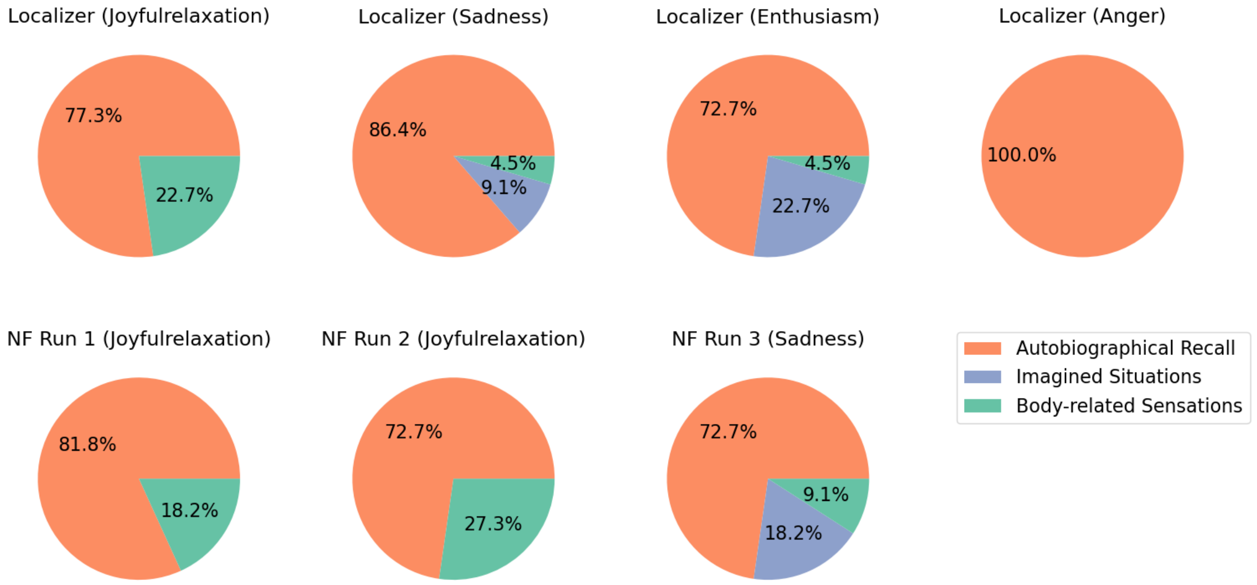


**Figure S8.** Distribution of emotion imagery strategies across participants. Questionnaires were collected for all localizer runs and the first two NF runs (N = 22), and for the final NF run (N = 11). Emotion imagination strategies were grouped into three categories: (1) Autobiographical recall — drawing on personal experiences (e.g., remembering a family gathering for joyful relaxation, or recalling a breakup to induce sadness); (2) Imagined situations — constructing hypothetical or unreal scenarios (e.g., picturing oneself winning a competition for enthusiasm, or being treated unfairly to elicit anger); and (3) Body-related sensations — focusing on physical feelings associated with emotions (e.g., a light and relaxed body state for joyful relaxation, or rising heat for anger). NF = neurofeedback.
